# Mettl3-catalyzed m^6^A methylation determines CD8^+^ T cell differentiation fate in tumor

**DOI:** 10.64898/2026.01.06.697843

**Authors:** Puspendu Ghosh, Anupam Gautam, Debashree Basak, Shaun Mahanti, Soham Chowdhury, Ishita Sarkar, Anwesha Mandal, Anwesha Kar, Snehanshu Chowdhury, Sandip Paul, Shilpak Chatterjee

## Abstract

The heterogeneity in patient responses to immune checkpoint blockade (ICB) is dictated by the relative abundance of exhausted CD8⁺ T cell (Tex) subsets with distinct therapeutic responsiveness. Progenitor exhausted (pTex) cells remain sensitive to ICB, whereas terminally exhausted (tTex) cells are refractory; however, the molecular cues that bias differentiation toward these divergent fates remain poorly defined. Here, we identify the RNA methyltransferase Mettl3 as a central regulator of Tex fate. Across murine tumor models, human T cells, and adoptive transfer systems, Mettl3 expression is selectively enriched in tTex cells and inversely correlated with TCF1⁺ pTex populations. Mechanistically, Mettl3 drives terminal exhaustion by stabilizing *DNMT3B* transcripts via m⁶A modification, enforcing CpG methylation and chromatin compaction at memory-associated loci. Inhibition of the Mettl3-Dnmt3b axis reprograms chromatin accessibility toward memory-like states, thereby preserving progenitor potential and effector function. Consequently, T cells lacking Mettl3-Dnmt3b activity persist longer, mount robust recall responses, and achieve superior tumor control with enhanced responsiveness to PD-1 blockade. These findings establish the Mettl3-m⁶A-Dnmt3b axis as a molecular rheostat of CD8⁺ T cell fate, coupling epitranscriptomic regulation to epigenetic remodeling, and reveal a tractable pathway to improve the durability of cancer immunotherapy.

## Introduction

The differentiation trajectory of naïve CD8⁺ T cells has been extensively characterized in the context of acute infection, where antigen-driven activation leads to robust clonal expansion and the generation of both short-lived effector cells and long-lived memory cells(*1*). In contrast to acute infections, the differentiation landscape of CD8⁺ T cells shifts dramatically during chronic infections and cancer(*2, 3*). Although these cells also undergo antigen-driven expansion, they fail to form functional memory populations and instead enter a dysfunctional state termed T cell exhaustion (Tex), driven by persistent antigen exposure(*2*). Tex cells exhibit sustained expression of multiple immune checkpoint molecules (PD-1, Tim-3, CD38, Lag-3), diminished effector activity, poor proliferative capacity, and impaired recall responses(*2, 4*). Current evidence largely supports a linear differentiation model for Tex development. In both chronic viral infections and cancer, at least two distinct Tex subsets have been identified: progenitor exhausted T cells (pTex) and terminally exhausted T cells (tTex), with pTex cells gradually giving rise to the more differentiated tTex population. pTex cells, maintained by the transcription factor TCF1, retain self-renewal capacity and are responsive to immune checkpoint blockade (ICB) therapy. In contrast, tTex cells, derived from pTex, lose TCF1 expression and become increasingly hyporesponsive to ICB(*5-11*). The considerable variability in patient responses to ICB therapy, even among those with similar disease profiles, challenges the strict linearity of the Tex differentiation model(*5, 12*). If the Tex program followed a truly linear trajectory, more consistent therapeutic responses would be expected. This discrepancy hints at the existence of alternative or parallel differentiation routes within the Tex compartment, contributing to functional and phenotypic heterogeneity and influencing therapeutic responsiveness.

Notably, a recent study demonstrated that intratumoral CD8⁺ T cells can acquire exhaustion-like features within just 24 hours of activation, challenging the conventional notion that dysfunction gradually accumulates through chronic stimulation(*13*). More intriguingly, this early fate commitment was accompanied by extensive changes in chromatin accessibility that established a stable program of dysfunction(*13*). These findings align with growing evidence that epigenetic remodeling is essential for Tex cell divergence from effector (Teff) and memory (T_Mem_) fates(*14, 15*). Once established, this altered chromatin landscape appears developmentally fixed, persisting even after antigen clearance. This raises an important and unresolved question: Are there specific cellular cues present during early CD8⁺ T cell activation that initiate epigenetic reprogramming and predispose cells toward a particular differentiation fate, be it exhaustion or memory? Identifying these signals will be key to stabilizing desirable CD8^+^ T cell states and improving immunotherapeutic efficacy.

Among the various regulatory layers, N6-methyladenosine (m^6^A), the most abundant internal modification in eukaryotic mRNA, has emerged as a key modulator of gene expression(*16, 17*). m^6^A is enriched near stop codons and within 3′ untranslated regions (3′ UTRs), and it influences mRNA stability, decay, and translation(*18-20*). This modification is deposited co-transcriptionally by the Mettl3-Mettl14 writer complex, with Mettl3 providing catalytic methyltransferase activity and Mettl14 contributing to RNA binding and complex stability(*21*). Mettl3-mediated m^6^A is critical for regulating cell fate and maintaining cellular homeostasis. In embryonic stem cells, for example, m^6^A facilitates differentiation by ensuring the timely downregulation of key naïve pluripotency-associated transcripts in response to developmental cues(*22*). Interestingly, genetic ablation of *Mettl3* in mouse T cells impedes their homeostatic proliferation and differentiation(*23*). Mettl3 is frequently overexpressed in several cancers, including oral and cutaneous squamous cell carcinoma, where it enhances stemness and drives tumor growth and invasiveness(*24-26*). Despite mounting evidence of Mettl3’s role across developmental contexts, its specific function in CD8⁺ T cell differentiation, particularly in cancer and the maintenance of exhausted (Tex) populations, remains poorly understood. Elucidating how Mettl3-mediated m^6^A intersects with epigenetic programming could uncover new strategies to modulate T cell fate for therapeutic benefit.

Here, we identify Mettl3 as a central determinant of CD8⁺ T cell fate in cancer. Through m⁶A-dependent stabilization of the epigenetic regulator Dnmt3b, Mettl3 enforces chromatin compaction at memory-associated loci, driving terminal exhaustion and suppressing progenitor potential. In contrast, inhibition of Mettl3 or Dnmt3b restores memory-like programs and markedly improves sensitivity to PD-1 blockade. These findings reveal an epitranscriptomic mechanism governing Tex differentiation and establish the Mettl3-m^6^A-Dnmt3b axis as a promising therapeutic target to enhance cancer immunotherapy.

## Results

### Mettl3 expression positively correlates with CD8^+^ T cell exhaustion

To investigate the role of Mettl3 in guiding the differentiation trajectory of exhausted CD8⁺ T cells (Tex), we employed two well-established murine tumor models, MC38 and YUMM1.7, which exhibit distinct intratumoral Tex profiles, specifically progenitor exhausted (pTex) versus terminally exhausted (tTex), and demonstrate differential responsiveness to immune checkpoint blockade (ICB) therapy(*27, 28*). We analyzed the temporal dynamics of T cell exhaustion and its relationship with Mettl3 expression by phenotyping CD8⁺ T cells from tumor tissues and tumor-draining lymph nodes (TdLN) of B6 mice subcutaneously implanted with either MC38 or YUMM1.7 tumors at multiple time points during tumor progression (Fig 1A). Consistent with prior reports, intratumoral antigen-experienced (CD44⁺) CD8⁺ T cells from both tumor types displayed canonical exhaustion markers, PD-1, Tim-3, and CD39, at higher levels than their TdLN counterparts (Extended Data Fig 1A). Further analysis revealed that YUMM1.7 tumors harbored a significantly higher frequency of tTex cells (PD1⁺Tim3⁺) compared to MC38 tumors, even at smaller tumor volumes (∼900 mm³). As tumor progression advanced, this enrichment of tTex in YUMM1.7 tumors became more pronounced. In line with this, PD1⁺CD39⁺ immunosuppressive Tex cells(*29*) were elevated in YUMM1.7 tumors both during early and late stages, whereas their prevalence remained comparatively lower in MC38 tumors (Fig S1A).

**Figure 1.**
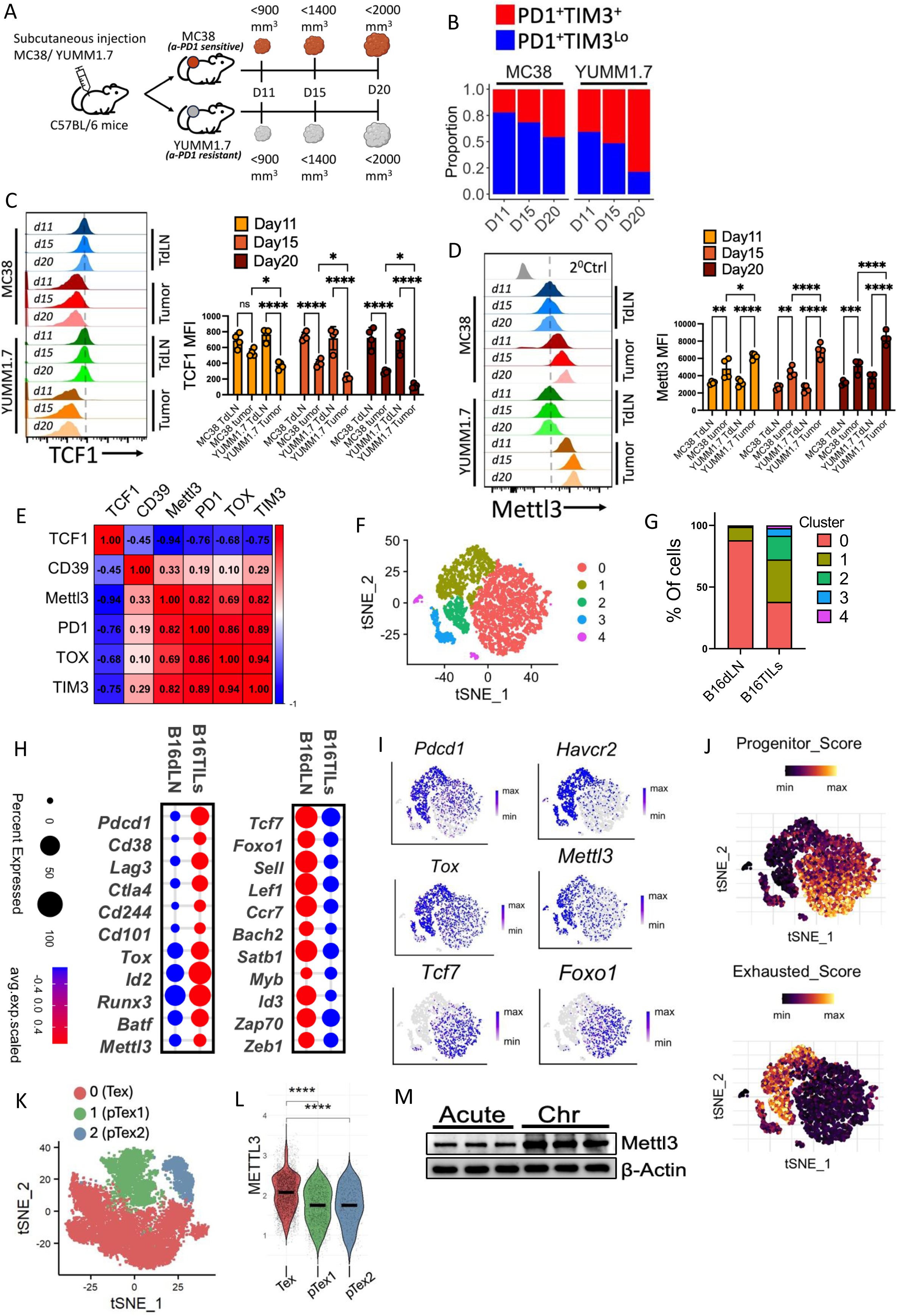
Expression of Mettl3 is primarily restricted to tTex cells. C57BL/6J mice were subcutaneously implanted with either MC38 or YUMM1.7 tumors, and intratumoral CD8⁺ T cells were analyzed at different time points. (A) Schematic representation of the experimental design. (B) proportion of PD1⁺Tim3⁺ (tTex) and PD1⁺Tim3^Lo^ (pTex) subsets in intratumoral CD8^+^T cells, (C) TCF1 expression, and (D) Mettl3 expression in CD8⁺ T cells during tumor progression. (E) Pearson correlation analysis of the indicated molecules in intratumoral CD8⁺ T cells isolated from MC38- and YUMM1.7-bearing mice at different time points. The adjacent bar diagrams (C & D) represent cumulative data from N=4 tumor-bearing mice. (F) tSNE visualization of the scRNA-seq clusters of murine CD8^+^ T cells from two samples. (G) Bar plot depicting the relative abundance of identified clusters across the two samples. (H) Dot plot representation of exhaustion-associated genes (left) and progenitor-associated genes (right) in CD8^+^ T cells. Dot size represents the percentage of cells expressing the gene, and color scale indicates average expression (scaled). (I) Single-cell transcription levels of representative genes illustrated in the tSNE plot. Transcription levels are color-coded: blue, expressed; gray, not expressed. (J) Module scores depicting enrichment of progenitor- and exhaustion-associated gene signatures. (K) tSNE projection of human scRNA-seq profiles from 4,588 intratumoral CD8⁺ T cells reveals three distinct CD8⁺ T cell subsets: Tex, pTex1, and pTex2. (L) Violin plot represents *METTL3* expression across three defined CD8⁺ T cell subsets. Significance was tested using two-sided Wilcoxon rank-sum with Bonferroni correction; (M) Mettl3 expression in CD8^+^ T cells from healthy human donors either stimulated acutely (Acute) or expanded with chronic antigen stimulation (Chr). The data are representative of three independent experiments. *, p < 0.05; **, p < 0.01; ***, p < 0.005; ****, p < 0.0001.

Conversely, the pTex population (PD1⁺Tim3^Lo^) remained relatively stable over time in MC38 tumors but declined markedly in YUMM1.7 tumors (Fig 1B). This trend was mirrored by the expression of TCF1, a transcription factor associated with the progenitor Tex pool: intratumoral CD8⁺ T cells from MC38 tumors maintained higher TCF1 expression throughout tumor progression than those from YUMM1.7 tumors (Fig 1C). Together, these findings suggest that the YUMM1.7 tumor microenvironment fosters a more profound and progressive state of T cell exhaustion compared to the MC38 model, potentially explaining its poorer response to ICB therapy.

Interestingly, longitudinal analysis of Mettl3 expression in intratumoral CD8⁺ T cells from both models revealed a reciprocal pattern to that of TCF1. Mettl3 levels were elevated from early time points in intratumoral CD8^+^ T cells from YUMM1.7 and continued to increase as the tumor progressed. In contrast, Mettl3 expression levels remained comparatively lower in CD8^+^T cells derived from the MC38 tumor site (Fig 1D). Correlation analysis further showed that Mettl3 expression positively correlated with key exhaustion markers, including PD1, Tim3, CD39, and TOX, but inversely correlated with TCF1, suggesting a mechanistic role for Mettl3 in driving terminal exhaustion at the expense of the progenitor pool (Fig 1E).

To investigate heterogeneity in Mettl3 expression across intratumoral CD8⁺ T cell subsets, we analyzed publicly available single-cell RNA-seq data of intratumoral CD8⁺ T cells from B16F10 melanoma-bearing mice(*27*). CD8⁺ T cells from the tumor-draining lymph node (TdLN) were also included to capture Mettl3 expression dynamics in tumor-specific stem-like memory T cells (T_SCM_) that are enriched in the TdLN(*30*). Integrating transcriptomic profiles from 9,056 CD8⁺ T cells across TME and TdLN revealed five distinct clusters (Fig 1F). Cluster 0, primarily contributed by TdLN-derived CD8⁺ T cells along with a fraction from the TME, exhibited high expression of memory-associated markers (*Tcf7, Foxo1, Sell, Lef1, Ccr7, Stab1,* and *Id3*) and was characterized as the pTex subset. In contrast, clusters 1 and 2, primarily contributed by intratumoral CD8^+^ T cells, labelled as Tex1 and Tex2, displayed elevated expression of multiple inhibitory receptors (*Pdcd1, Havcr2, Lag3, Ctla4,* and *Cd38*) along with exhaustion-associated transcription factors (*Tox* and *Id2*), but lacked *Tcf7* expression (Fig 1G-1H). Average scaled expression of *Mettl3* revealed that it was predominantly enriched in clusters 1 and 2, whereas it was largely absent or expressed at minimal levels in cluster 0 (Fig 1I). This distribution suggests that Mettl3 expression is more closely associated with tTex cells than with the pTex subset, as defined by previously established gene signatures(*11*) for these populations (Fig 1J). To assess whether this pattern is conserved in human tumors, we analyzed publicly available scRNA-seq datasets (derived from 10X Genomics Datasets) from five different malignancies: breast cancer (two independent datasets), colorectal cancer, endocervical adenocarcinoma, and skin melanoma. After stringent quality control and filtering, unsupervised clustering of the integrated datasets identified 14 clusters (Fig S1B). Cluster 5 was annotated as intratumoral CD8⁺ T cells based on the expression of lineage-defining markers *CD8A* and *CD3G* (Fig S1C). A t-SNE projection of 4588 CD8⁺ T cells from cluster 5 revealed three distinct clusters (Fig 1K). Cluster 0 corresponded to Tex cells, defined by high expression of exhaustion-associated genes (*PDCD1, TIGIT, TOX, CD244*) and elevated exhaustion module scores (Fig S1D-S1E). In contrast, clusters 1 and 2, labeled pTex1 and pTex2, were enriched for memory-associated genes. pTex1 expressed *FOXO1, CXCR5, IL2RB, and CD28*, while pTex2 expressed *SELL, TCF7, LEF1,* and *SLAMF6*; both subsets displayed high progenitor module scores (Fig S1D-S1E). Strikingly, *METTL3* expression was significantly higher in Tex cells (cluster 0) compared with both progenitor clusters (1 and 2) (Fig 1L). These findings mirror our observations in murine tumors, reinforcing the preferential association of Mettl3 with the terminally exhausted subset and its inverse relationship with progenitor subsets.

Finally, we assessed Mettl3 expression in chronically stimulated human CD8⁺ T cells to determine whether its elevated levels in Tex cells result from chronic TCR stimulation. For this, CD8⁺ T cells from healthy human donors were chronically stimulated to induce exhaustion, which was confirmed by an increased frequency of PD1⁺Tim3⁺ cells (Fig. S1F), elevated CTLA4 expression (Fig. S1G), reduced TCF1 levels (Extended Data Fig. 1H), and functional impairment, as evidenced by diminished IFNγ and TNFα production (Fig. S1I-S1J). Notably, these chronically exhausted CD8⁺ T cells exhibited markedly higher Mettl3 expression compared with acutely stimulated CD8⁺ T cells (Fig. 1L).

Collectively, these findings indicate that Mettl3 expression is preferentially enriched in terminally exhausted CD8⁺ T cells and inversely associated with the progenitor pool.

### Mettl3 inhibition alleviates CD8^+^ T cell exhaustion while promoting T_Mem_ generation in vitro

Given the reciprocal association of Mettl3 with tTex and pTex subsets, we sought to determine whether Mettl3 expression differentially regulates the differentiation of Tex and memory-like subsets following CD8⁺ T cell activation. To investigate this, human CD8⁺ T cells purified from healthy donors were activated for 3 days and then cultured for an additional 12 days under either chronic antigen stimulation with plate-bound anti-CD3 to promote Tex differentiation or with IL-15 and IL-7 supplementation, in the absence of antigen, to favor T_Mem_ differentiation (Fig 2A). Mettl3 expression was monitored at multiple time points up to day 15 to assess its temporal dynamics during differentiation. Interestingly, CD8⁺ T cells rapidly upregulated Mettl3 upon activation (Fig S2A), and its expression remained elevated and progressively increased under chronic TCR stimulation (Fig 2B and 2D), suggesting a link with Tex differentiation. In contrast, under memory-inducing conditions (IL-15/IL-7), Mettl3 expression gradually declined over time, consistent with a reduced role in T_Mem_ generation (Fig 2C and 2D).

**Figure 2.**
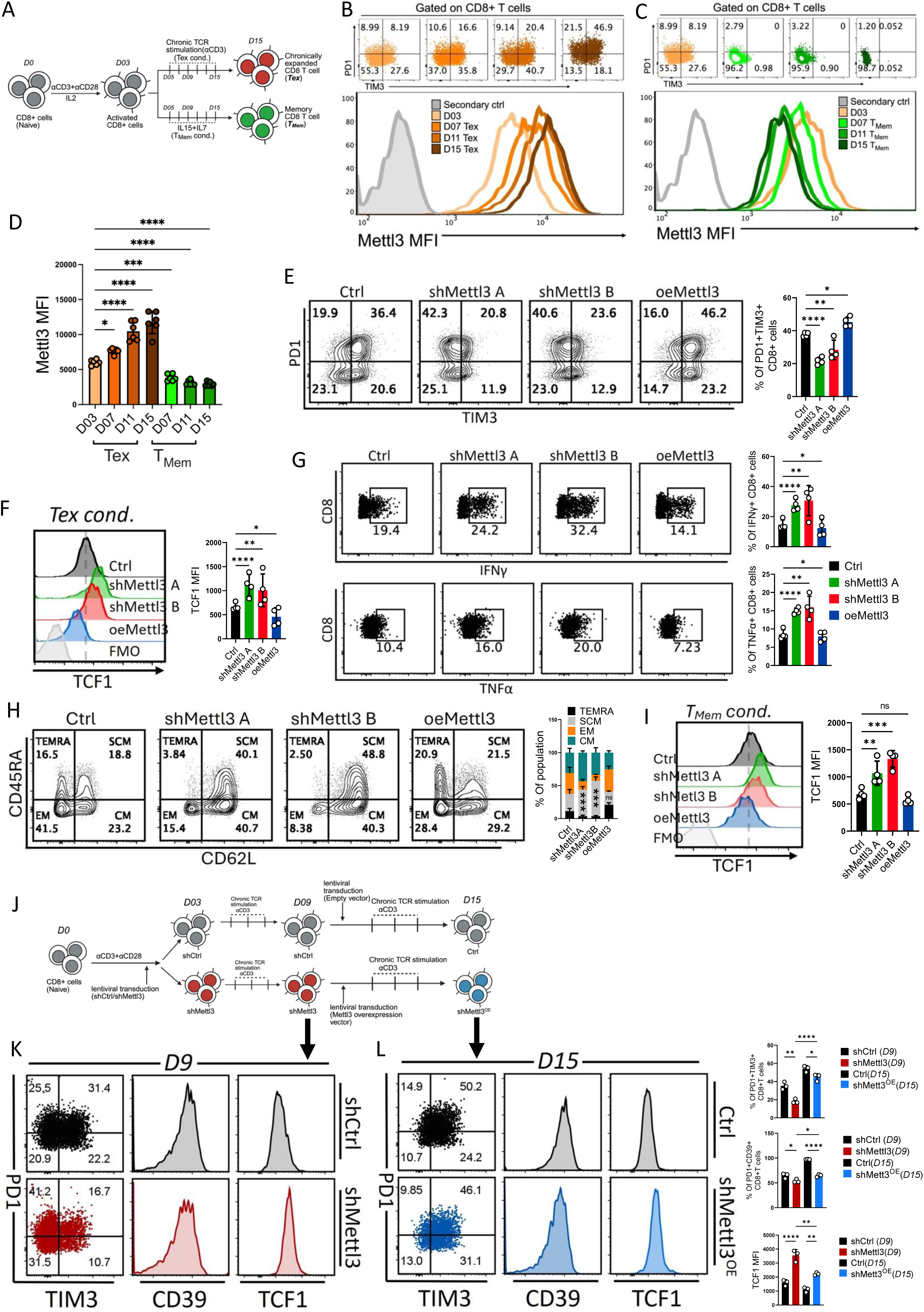
Mettl3 expression promotes exhaustion while impairing memory-like CD8⁺ T cell differentiation. (A) Schematic of in vitro differentiation of Tex and T_Mem_ cells from healthy human donors. (B-D) Temporal dynamics of Mettl3 expression in in vitro differentiated (B) Tex cells and (C) T_Mem_ cells. (D) Bar graph summarizing cumulative data from six independent experiments. (E-G) Activated CD8^+^ T cells were transduced with either two independent shRNA constructs targeting Mettl3 (shMettl3 A and shMettl3 B) or a Mettl3 overexpression vector (oeMettl3), chronically expanded up to D15 and analyzed for: (E) PD1 and Tim3 expression, (F) intracellular TCF1 levels, and (G) cytokine production. (H-I) In vitro-differentiated T_Mem_ cells from the indicated groups were analyzed for: (H) CD45RA and CD62L expression, and (I) TCF1 expression. Adjacent bars show cumulative data from four independent experiments for (E-I). (J) Schematic of the experimental design. (K) At D9, CD8⁺ T cells transduced with either control (Ctrl) or Mettl3 shRNA (shMettl3) were analyzed for: PD1 and Tim3, CD39, and TCF1 expression. (L) At D15, CD8⁺ T cells from (K) overexpressing Mettl3 (shMettl3^OE^) were assessed for: PD1 and Tim3, CD39, and TCF1 expression. Adjacent bar graphs show combined data from D9 and D15 samples (K-L), pooled from three independent experiments. *, p < 0.05; **, p < 0.01; ***, p < 0.005; ****, p < 0.0001.

To delineate the functional role of Mettl3, we performed both genetic knockdown and overexpression of *METTL3* in human CD8⁺ T cells (Fig. S2B) and subjected them to either Tex or T_Mem_ differentiation. Loss of Mettl3 markedly impaired Tex differentiation under sustained TCR stimulation, as indicated by reduced frequencies of CD8⁺ T cells co-expressing canonical exhaustion markers (PD1⁺Tim3⁺, PD1⁺CD39⁺, and PD1⁺CTLA4⁺), along with a notable increase in TCF1 expression (Fig 2E-2F, and Fig S2C). In contrast, Mettl3 overexpression enhanced T cell exhaustion, reflected by higher frequencies of PD1⁺Tim3⁺, PD1⁺CD39⁺, and PD1⁺CTLA4⁺ CD8⁺ T cells and a concomitant reduction in TCF1 levels (Fig 2E-2F, and Fig S2C). Functionally, Tex cells generated under Mettl3-deficient conditions exhibited improved cytokine production (IFN-γ and TNF-α), whereas Mettl3 overexpression had minimal effects on cytokine output compared with controls (Fig 2G).

In sharp contrast, Mettl3 knockdown under memory-differentiation conditions promoted the formation of T_Mem_ cells, particularly the long-lived, multipotent stem cell memory (SCM)-like subset characterized by co-expression of CD45RA and CD62L. In contrast, overexpression of Mettl3 led to only a modest increase in the TEMRA population and failed to promote differentiation of the SCM-like subset (Fig 2H). Importantly, Mettl3-deficient cells exhibited elevated expression of memory-associated markers, including TCF1, FOXO1, and CCR7, and mounted stronger recall responses, as evidenced by enhanced effector cytokine production upon re-stimulation (Fig 2I and Fig S2D-S2F).

To further confirm whether Mettl3 directly governs the fate decision between exhausted and memory-like CD8⁺ T cell states, we performed a rescue experiment. CD8⁺ T cells were first subjected to Mettl3 knockdown and exposed to chronic stimulation to induce exhaustion. Mettl3 was subsequently reintroduced into the same cells, which were then maintained under identical Tex-promoting conditions (Fig 2J). As expected, Mettl3 knockdown reduced the frequency of exhausted CD8⁺ T cells (PD1⁺Tim3⁺ and CD39⁺) while increasing TCF1 expression. In contrast, reintroduction of Mettl3 restored the exhausted phenotype in chronically stimulated CD8⁺ T cells, as reflected by an increased frequency of PD1⁺Tim3⁺, and CD39⁺ cells, accompanied by reduced TCF1 expression (Fig 2K-2L). Together, these findings demonstrate that Mettl3 actively promotes CD8⁺ T cell exhaustion while restraining memory-like differentiation.

Collectively, these findings identify Mettl3 as a pivotal regulator of CD8⁺ T cell fate. Depending on its temporal expression dynamics, Mettl3 orchestrates the balance between terminal exhaustion and long-lived memory differentiation, likely by modulating the underlying molecular circuitry that governs lineage commitment.

### Mettl3-induced epigenetic bias limits T_Mem_ differentiation in vitro

Previous studies have demonstrated that during acute infection, CD8⁺ T cells commit to an ‘instructive developmental program’ early during activation, which governs their differentiation into effector and memory subsets(*31, 32*). Given the observed role of Mettl3 in promoting distinct CD8⁺ T cell fates, favoring Tex over T_Mem_, we next investigated whether Mettl3 activity during the early activation phase programs the subsequent differentiation trajectory of CD8⁺ T cells. To selectively inhibit catalytic function of Mettl3 during activation, without inducing stable gene knockdown, we employed STM2457 (hereafter referred to as STM), a pharmacological inhibitor that specifically targets the methyltransferase domain of Mettl3. To validate the efficacy of STM, human CD8⁺ T cells were activated in the presence of STM (10 μM), and RNA was extracted on day 3 to assess global m^6^A methylation. STM-treated cells showed a marked reduction in m^6^A levels compared to vehicle-treated controls, confirming effective inhibition of Mettl3 enzymatic activity (Fig S3A). Importantly, STM treatment did not compromise overall T cell activation (Fig S3B-S3C).

Next, purified human CD8⁺ T cells were activated in the presence or absence of STM for three days. After activation, cells were thoroughly washed to remove residual inhibitor and then cultured under Tex- or T_Mem_-polarizing conditions in inhibitor-free media, as outlined schematically (Fig 3A). Interestingly, transient inhibition of Mettl3 during activation hindered terminal differentiation and promoted the retention of a progenitor-like phenotype. This was reflected by a reduced frequency of CD8⁺ T cells co-expressing PD1⁺Tim3⁺ and PD1⁺CD39⁺, hallmarks of terminal exhaustion, while sustaining TCF1 expression even under Tex-inducing conditions (Fig 3B-3D). Functionally, STM-treated CD8⁺ T cells also exhibited enhanced cytokine production compared to untreated cells (Fig 3E-3F).

**Figure 3.**
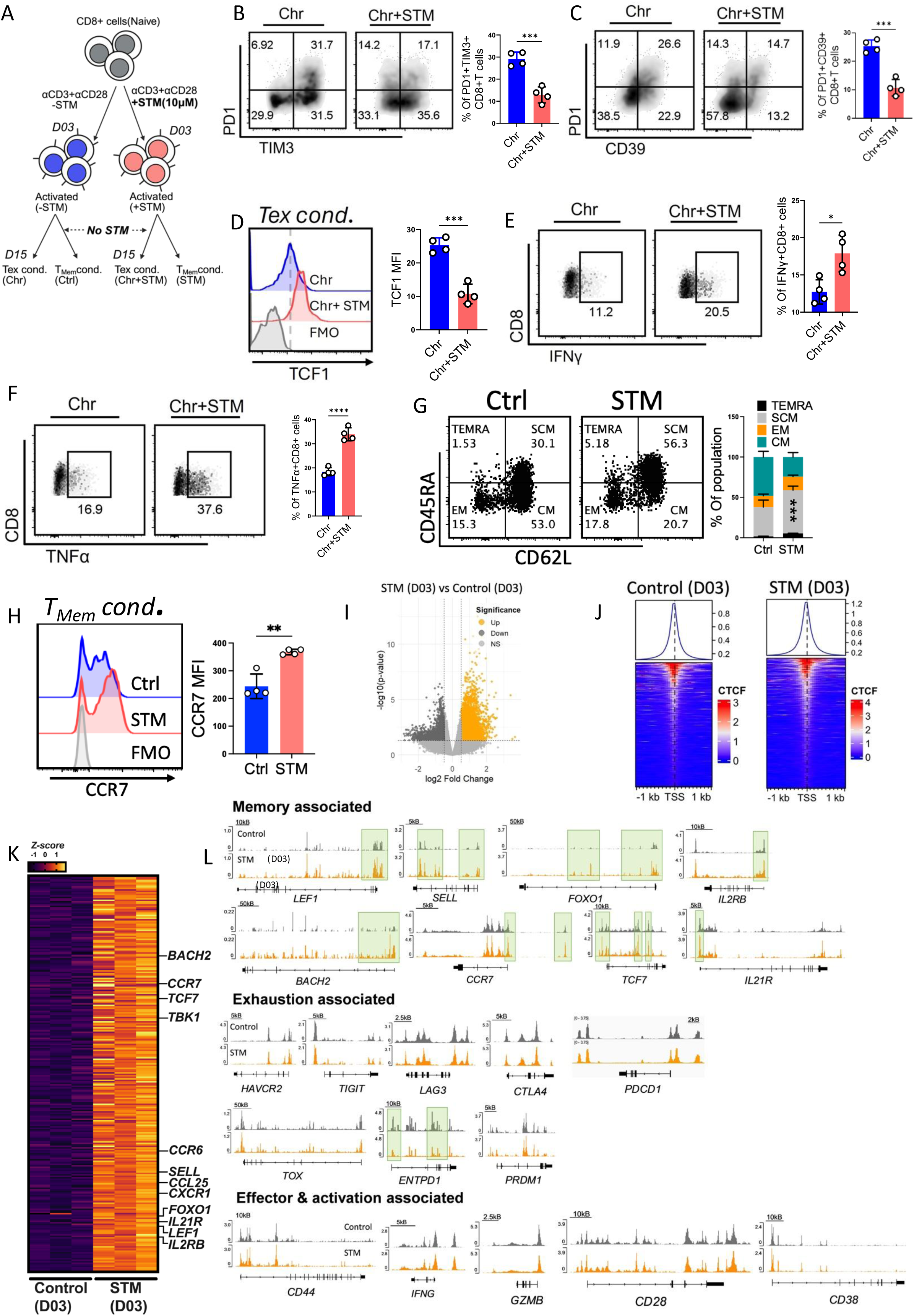
Mettl3 confers an epigenetic bias that limits T_Mem_ generation. (A) Schematic of the experimental design: CD8⁺ T cells from healthy human donors were activated in the presence or absence of STM and then subjected to either Tex differentiation conditions (Tex Cond.) or T_Mem_ differentiation conditions (T_Mem_ Cond.) in the absence of STM. (B-F) CD8⁺ T cells cultured under Tex Cond., as in (A), were analyzed for: (B) PD1 and TIM3 expression, (C) PD1 and CD39 expression, (D) TCF1 expression, (E) intracellular IFNγ production, and (F) intracellular TNFα production. Adjacent bar graphs (B-F) represent cumulative data from four independent experiments. (G, H) CD8⁺ T cells cultured under T_Mem_ Cond., as in (A), were analyzed for the expression of: (G) CD45RA and CD62L, and (H) CCR7. The adjacent bar graph (G), representative of four independent experiments, shows the frequency of memory-associated subsets defined by CD45RA and CD62L expression, and (H) shows cumulative data from four independent experiments. (I) Volcano plot of 158,435 differentially accessible chromatin regions (DACRs) identified by DESeq2. (J) CTCF binding profiles at transcription start sites (±1 kb) under Control and STM conditions. Line graphs represent average signal intensity; heatmaps display locus-specific binding strength from low (blue) to high (red). (K) Heat map of the top 250 differentially accessible promoter regions (TSS) between control and STM-treated cells. (L) ATAC-seq tracks showing chromatin accessibility of memory-, exhaustion-, and effector/activation-associated genes in control versus STM-treated CD8⁺ T cells. Peaks uniquely lost or gained are highlighted in green. *, p < 0.05; **, p < 0.01; ***, p < 0.005; ****, p < 0.0001.

In contrast, under T_Mem_-inducing conditions, Mettl3 inhibition during activation favored the generation of a stem cell memory (SCM)-like population (Fig 3G), recapitulating the phenotype observed with Mettl3 knockdown. This was accompanied by upregulation of memory-associated surface markers CCR7 (Fig 3H), indicating a preferential bias toward T_Mem_ differentiation imprinted during the early activation window.

Given that enhanced mitochondrial function is a hallmark of long-lived memory T cells(*33, 34*), we next profiled the metabolic features of CD8⁺ T cells activated in the presence or absence of STM. STM-treated cells exhibited a marked reduction in glycolytic activity, consistent with the downregulation of key glycolysis-related genes, likely due to m^6^A-dependent regulation of transcript stability, as previously reported(*35*) (Extended Data Fig 3D-3E). In contrast, mitochondrial oxidative phosphorylation (OXPHOS), particularly the spare respiratory capacity (SRC), a metabolic prerequisite for memory formation, was significantly elevated in STM-treated cells (Fig S3F). In line with this, both mitochondrial abundance and activity were increased, as indicated by elevated Mitogreen and Mitored staining, respectively (Fig S3G-S3H). Furthermore, expression of the mitochondrial fusion-related gene *Opa1* was significantly upregulated in STM-treated CD8⁺ T cells (Fig S3I).

Memory-precursor and stem-like T cells have been reported to show reduced proliferation despite active differentiation(*36*). Using Cell Trace Violet (CTV) dilution, we observed that while the total number of cell divisions remained comparable between STM-treated and control groups, the frequency of cells in later divisions (P4+P5) was substantially lower in the STM group (Fig S3J). BrdU incorporation further revealed that a significant fraction of STM-treated cells were in the G2/M phase, suggesting reduced, but not arrested, proliferative activity (Fig S3K). Together, these findings suggest that inhibition of Mettl3 during early activation alters both metabolic and proliferative dynamics to favor memory-like fate acquisition.

To determine whether this memory-prone differentiation bias was epigenetically encoded during activation, we performed transposase-accessible chromatin using sequencing (ATAC-seq) on CD8⁺ T cells activated in the presence or absence of STM. Differential analysis revealed 158,435 differentially accessible chromatin regions (DACRs) between the two groups, with a prominent enrichment of accessibility around transcription start sites (TSS, −1 to +1 kb) (Fig 3I and 3J). Notably, STM-treated CD8⁺ T cells exhibited increased chromatin accessibility at the promoter-TSS regions of several key memory-associated genes, including *LEF1*, *SELL*, *FOXO1*, *IL2RB*, *BACH2*, *CCR7*, *TCF7*, and *IL21R* (Fig 3K-3L). In contrast, the accessibility at exhaustion-related loci (*HAVCR2*, *TIGIT*, *LAG3*, *CTLA4*, *PDCD1*, *TOX*, *PRDM1*), with the exception of *ENTPD1*, remained largely unaltered. Similarly, genes involved in effector function or activation (*CD44*, *IFNG*, *GZMB*, *CD28*, *CD38*) showed no significant differences in accessibility between groups (Fig 3L).

Together, these findings reveal that Mettl3 inhibition during early CD8⁺ T cell activation selectively remodels the chromatin landscape to favor a memory-like transcriptional program. This early epigenetic imprinting likely underpins the long-lived, multipotent phenotype observed upon transient loss of Mettl3 activity.

### m^6^A-Dependent stabilization of Dnmt3b by Mettl3 Orchestrates Epigenetic Remodeling

Mettl3-catalyzed m⁶A is the most prevalent epitranscriptomic modification, regulating mRNA stability, translation, splicing, and nuclear export. Pharmacological inhibition of Mettl3 methyltransferase activity with STM altered the chromatin landscape of CD8⁺ T cells, prompting us to investigate the underlying molecular mechanism. We performed global RNA sequencing (RNA-seq) on human CD8⁺ T cells activated with or without STM. Comparative analysis identified 1,185 differentially expressed genes (DEGs; 550 upregulated, 635 downregulated; log₂FC ≥ 0.3, padj < 0.05). Gene Ontology analysis (GO) revealed that STM-treated CD8⁺ T cells were enriched for pathways associated with long-lived memory T cell development, including DNA damage response, Wnt signaling, mitochondrial gene expression, and stem cell proliferation (Fig 4A). Further inspection of DEGs showed elevated expression of memory-associated genes (*LEF1, CCR7, FOXO1, BCL11B, BCL2,* and *IL6R*), whereas genes implicated in negative regulation of T cell function (*ENTPD1, HAVCR2, TNFRSF8, TNFRSF18*, and *CTLA4*) were downregulated (Fig 4B). Interestingly, STM treatment also increased TRAF6 expression (Fig 4B), which is known to promote CTLA4 ubiquitination and degradation(*37*).

**Figure 4.**
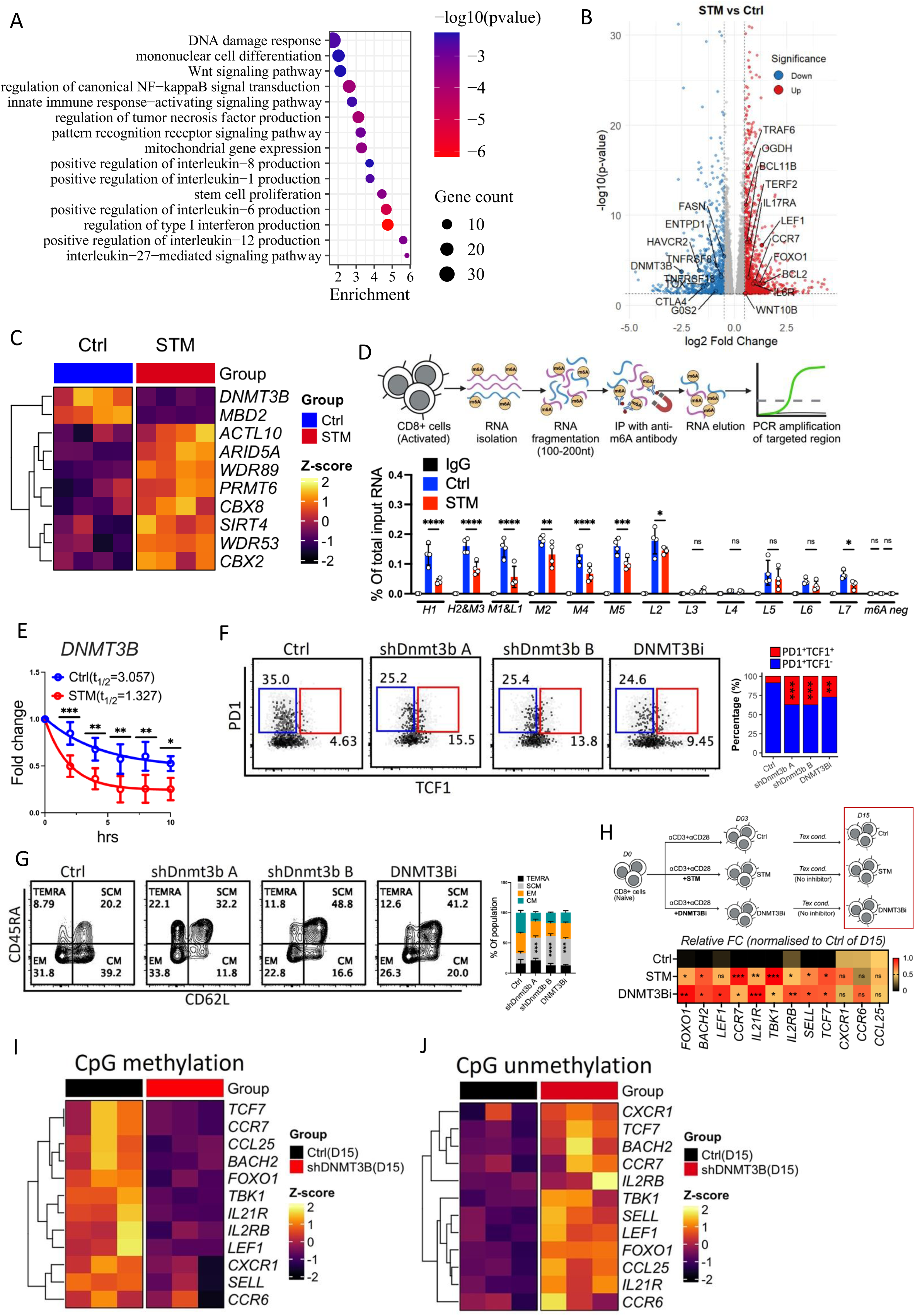
The Mettl3-m⁶A-Dnmt3b axis promotes epigenetic remodeling at memory-associated genes. (A, B) CD8⁺ T cells from healthy human donors were activated for three days in the presence or absence of STM and subjected to bulk RNA-seq analysis. (A) Top 15 GO pathways enriched in STM-treated CD8⁺ T cells compared to control. (B) Volcano plot showing differentially expressed genes in STM vs Ctrl CD8⁺ T cells. Significant genes are colored; gray dots indicate non-significant changes; selected genes are labeled. (C) Heat map showing differentially expressed epigenetic modulators. (D) Schematic of the m⁶A-RIP workflow (upper panel). Bar plots (bottom panel) show enrichment of m⁶A-modified regions in *DNMT3B* transcripts. (E) *DNMT3B* mRNA decay in control and STM-treated CD8⁺ T cells following Actinomycin D treatment. Decay rates were determined by non-linear regression curve fitting (one-phase decay model). Data are representative of four independent experiments. (F) PD1 versus TCF1 expression in chronically exhausted CD8⁺ T cells upon Dnmt3b knockdown (shDnmt3b A and shDnmt3b B) or pharmacological inhibition (DNMT3Bi). The adjacent bar graph shows cumulative frequencies of PD1⁺TCF1⁺ and PD1⁺TCF1⁻ subsets from four independent experiments. (G) Memory subset distribution based on CD45RA and CD62L expression in CD8⁺ T cells differentiated under T_Mem_ conditions. The adjacent bar plot shows the percentage of each memory subset from four independent experiments. (H) Schematic of experimental design (upper panel); qPCR analysis of key memory-associated transcripts (bottom panel). Data are representative of four independent experiments. (I, J) Heat maps showing (I) CpG methylation and (J) unmethylation status at key memory-associated gene loci following bisulfite conversion and PCR amplification using methylation- and unmethylation-specific primers, respectively. Data are representative of three independent experiments. *, p < 0.05; **, p < 0.01; ***, p < 0.005; ****, p < 0.0001, ns, non-significant.

To explore the link between Mettl3 inhibition and epigenetic remodeling, we analyzed DEGs involved in epigenetic regulation. Among various modifiers, *DNMT3B*, encoding the de novo DNA methyltransferase Dnmt3b, was markedly downregulated in STM-treated CD8⁺ T cells (Fig 4C). qPCR-based RT-profiler array validation further confirmed *DNMT3B* as the most reduced transcript (∼2-fold) among all tested modifiers (Fig S4A).

Because m⁶A modification by Mettl3 regulates mRNA stability, we next examined whether *DNMT3B* downregulation stemmed from reduced transcript stability. SRAMP database analysis predicted 14 potential m⁶A sites on *DNMT3B* mRNA with high, moderate, or low confidence scores (Fig S4B-S4C). RNA immunoprecipitation (RNA-IP) with anti-m⁶A antibody in STM-treated and control CD8⁺ T cells, followed by qPCR of predicted regions (a non-m⁶A region as controls), confirmed m⁶A modification at most high/moderate sites in *DNMT3B* transcript, which was substantially reduced by STM treatment (Fig 4D). Furthermore, Actinomycin D chase assay unveiled a marked decline in the half-life (t_₁/₂_) of *DNMT3B* mRNA following Mettl3 inhibition, underscoring the critical role of Mettl3-mediated m⁶A methylation in preserving the stability of *DNMT3B* transcripts (Fig 4E). This effect was specific to *DNMT3B*, as *DNMT3A* and *DNMT1* stability remained unaffected upon STM treatment (Fig S4D).

To assess whether Mettl3-mediated regulation of CD8⁺ T cell differentiation involves the epigenetic modifier Dnmt3b, we first examined its expression in intratumoral CD8⁺ T cells from MC38 and YUMM1.7 tumors. Similar to Mettl3, Dnmt3b expression was higher in YUMM1.7-derived intratumoral CD8⁺ T cells compared to those from MC38 tumors (Fig S4E). We next evaluated the functional role of Dnmt3b. Similar to Mettl3, Dnmt3b expression progressively increased as CD8⁺ T cells underwent chronic stimulation and differentiated into Tex but displayed the opposite trend during in vitro differentiation into T_Mem_ (Fig S4F). Genetic knockdown of *DNMT3B* or catalytic inhibition using a pharmacological inhibitor (Nanaomycin A, hereafter referred to as DNMT3Bi) in human CD8⁺ T cells had only modest effects on exhaustion marker expression during Tex differentiation but stabilized the progenitor-like subset, as indicated by increased TCF1 expression with heightened effector function (Fig 4F and Fig S4G-S4I). Notably, under T_Mem_ differentiation conditions, Dnmt3b inhibition or knockdown enriched the stem cell-like memory (SCM; CD45RA⁺CD62L⁺) population, upregulated memory-associated transcription factors (FOXO1 and TCF1) and the homing marker CCR7, and enhanced effector function upon restimulation. (Fig 4G and Fig S4J-S4M). Together, these findings suggest that Dnmt3b activity is essential for regulating memory-associated genes but largely dispensable for maintaining exhaustion-related programs.

Since Dnmt3b-mediated CpG methylation promotes chromatin closure and represses transcription, we investigated whether elevated Dnmt3b in Tex cells silences genes associated with memory and multipotency. Tex cells treated with DNMT3Bi exhibited marked upregulation of long-lived memory–associated genes, including *FOXO1, BACH2, LEF1, CCR7, IL21R, TBK1, IL2RB, SELL,* and *TCF7* (Fig 4H). Consistently, similar upregulation of these genes was also observed upon inhibition of Mettl3 activity using STM, supporting our finding that Mettl3 functions as an upstream regulator of Dnmt3b (Fig 4H). To further examine whether this regulation is driven by Dnmt3b-mediated CpG methylation at the promoter, we performed bisulfite conversion of genomic DNA and analyzed CpG methylation indetified within the 4 kb upstream of transcriptional start site (TSS) of these genes in CD8⁺ T cells transduced with control or *DNMT3B* shRNA during Tex differentiation. Fifteen days post-differentiation, memory-associated genes in chronically exhausted CD8^+^ T cells (Ctrl) were heavily methylated, as evidenced by amplification with methylation-specific but not unmethylation-specific primers (Fig 4I-4J). In contrast, *DNMT3B* knockdown (shDNMT3B) reversed this epigenetic silencing, leading to unmethylated CpGs and subsequent amplification with unmethylation-specific primers, indicating restoration of a more permissive chromatin state (Fig 4I-J). These findings confirm that Dnmt3b-mediated methylation restricts promoter accessibility of memory-associated genes, thereby limiting the differentiation potential of pTex cells while facilitating terminal differentiation. Notably, these closed chromatin states, marked by high methylation, emerged in CD8⁺ T cells immediately after activation, suggesting that Dnmt3b activity transcriptionally represses memory-associated genes in effector CD8⁺ T cells. However, inhibition of Dnmt3b during activation epigenetically predisposed the cells to adopt a memory fate by maintaining open chromatin at memory-associated gene loci.

### Targeting Mettl3 or Dnmt3b improves the memory potential and anti-tumor efficacy of CD8^+^ T cells in vivo

Having established that the Mettl3-Dnmt3b axis limits the expression of long-lived memory-associated genes in CD8⁺ T cells by modulating chromatin accessibility, we next investigated whether targeting either Mettl3 or Dnmt3b could enhance CD8⁺ T-cell persistence and elicit a stronger anti-tumor immune response. To address this, we performed the T cell parking experiment in Rag1^-/-^ mice. Pmel T cells-harboring a transgenic TCR specific for the gp100 antigen expressed by B16F10 melanoma-were transduced with either control shRNA (Pmel^Wt^), Mettl3 shRNA (Pmel^shMettl3^) or Dnmt3b shRNA (Pmel^shDnmt3b^), activated for three days with the cognate peptide, and adoptively transferred into Rag1^⁻/⁻^ mice without tumor engraftment, as outlined schematically (Fig 5A). Prior to adoptive transfer, knockdown efficiency was confirmed by assessing Mettl3 and Dnmt3b expression in Pmel^shMettl3^ and Pmel^shDnmt3b^ cells, respectively (Fig S5A-S5B). Twenty-one days post-transfer, peripheral blood was analyzed for persistence and memory-associated markers in the transferred TCR transgenic Pmel T cells. Our analysis revealed that Pmel^shMettl3^ and Pmel^shDnmt3b^ cells persisted at higher frequencies and displayed a memory phenotype, as evidenced by increased central memory (CD44⁺CD62L⁺) populations and elevated expression of CCR7, TCF1, and FOXO1 (Fig 5B and Fig S5C-S5F).

**Figure 5.**
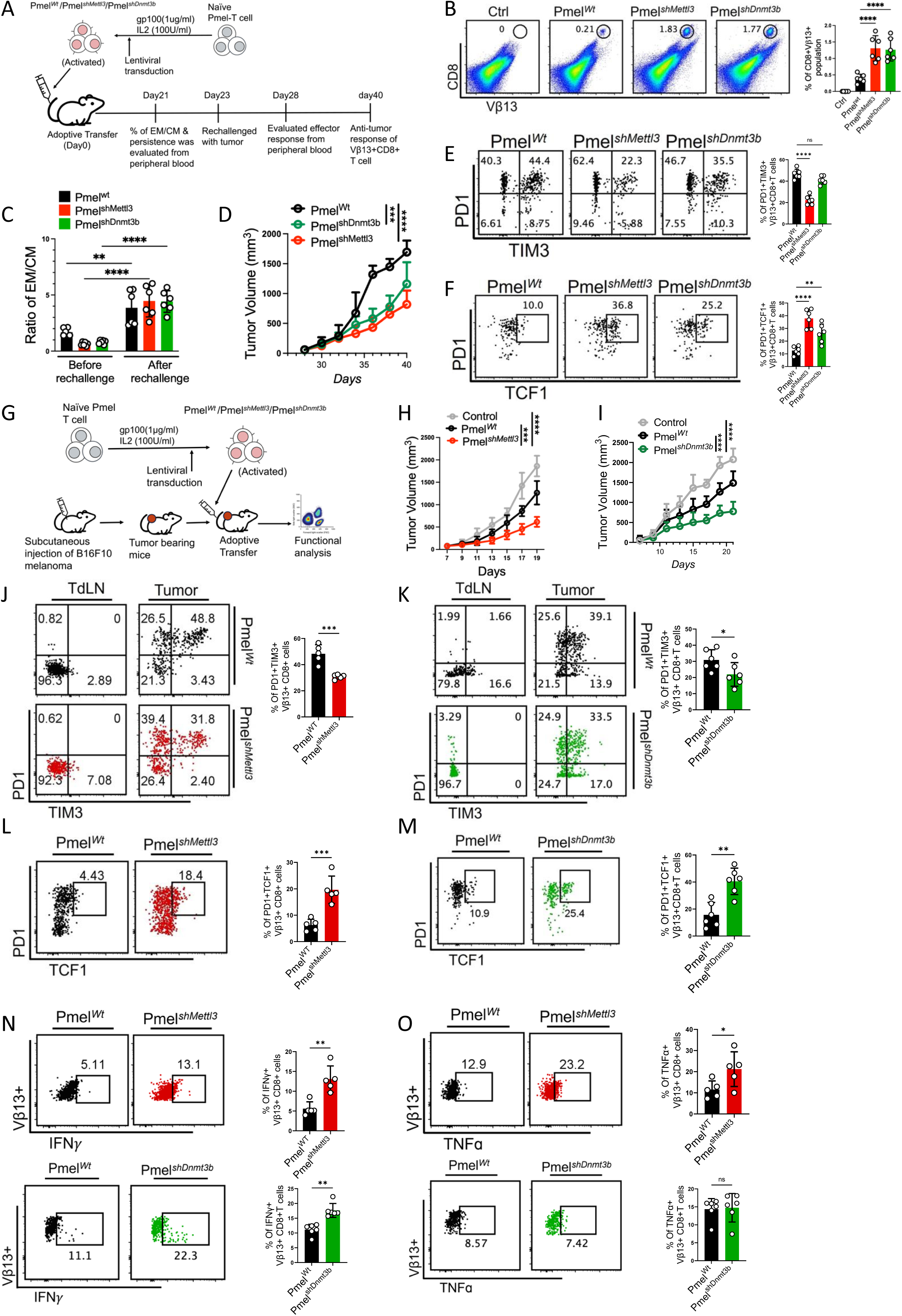
Genetic knockdown of Mettl3 or Dnmt3b enhances CD8⁺ T cell persistence and anti-tumor activity. (A) Schematic of the experimental design: activated Pmel T cells transduced with either control shRNA or Mettl3 shRNA or Dnmt3b shRNA were adoptively transferred (1.5 × 10⁶ cells/mouse) into Rag1⁻/⁻ mice (N = 6 mice/group). (B) Frequency of Pmel TCR transgenic T cells (CD8⁺Vβ13⁺) in peripheral blood at day 21 post-transfer. On day 23, mice were challenged with B16F10 melanoma by subcutaneous implantation, and on day 28, peripheral blood was analyzed for (C) the ratio of effector memory (EM; CD44⁺CD62L⁻) to central memory (CM; CD44⁺CD62L⁺) subsets within CD8⁺Vβ13⁺ T cells, compared with the profile obtained on day 21 (before rechallenge). (D) Tumor growth kinetics in the same cohorts. Intratumoral CD8⁺Vβ13⁺ T cells were further analyzed for (E) PD1 and Tim3 expression and (F) PD1 and TCF1 expression. (G-O) Rag1⁻/⁻ B6 mice (N = 6 mice/group) bearing subcutaneous B16F10 tumors were adoptively transferred with either wild-type Pmel (Pmel^Wt^), Mettl3-knockdown Pmel (Pmel^shMettl3^), or Dnmt3b-knockdown Pmel (Pmel^shDnmt3b^) T cells (1.5 × 10⁶ cells/mouse). (G) Schematic of the experimental design. Tumor-bearing mice were assessed for: (H, I) tumor volume; (J, K) frequency of CD8⁺Vβ13⁺ T cells expressing PD1 and Tim3; (L, M) frequency of CD8⁺Vβ13⁺ T cells expressing PD1 and TCF1; and (N, O) intracellular IFNγ and TNFα production. Adjacent bars represent cumulative data from N = 6 mice. *, p < 0.05; **, p < 0.01; ***, p < 0.005; ****, p < 0.0001.

To assess recall capacity, a hallmark of functional memory T cells, the same recipient mice were challenged with B16F10 melanoma via subcutaneous implantation. Five days later, peripheral blood was analyzed again. Both Pmel^shMettl3^ and Pmel^shDnmt3b^ cells, in response to antigen re-encounter, transitioned more efficiently to effector memory (CD44⁺CD62L⁻) subsets compared to Pmel^Wt^ cells (Fig 5C and Extended Data Fig 5G). Notably, even after antigen re-exposure, Pmel^shMettl3^ and Pmel^shDnmt3b^ retained significantly higher expression of CCR7, TCF1, and FOXO1, indicating preservation of a progenitor memory pool with self-renewal and persistence potential (Fig S5H-S5J). Consistent with these findings, tumor growth monitoring revealed that Rag1^⁻/⁻^ mice receiving either Pmel^shMettl3^ or Pmel^shDnmt3b^ cells mounted a more potent anti-tumor response than those receiving Pmel^Wt^ cells (Fig 5D). Analysis of Vβ13⁺ T cells (the transgenic TCR expressed by Pmel T cells) from tumors showed that both Pmel^shMettl3^ and Pmel^shDnmt3b^ T cells exhibited reduced features of terminal exhaustion, including lower frequencies of PD1⁺Tim3⁺, PD1⁺CTLA4⁺, and PD1⁺CD39⁺ cells, compared with Pmel^Wt^ cells (Fig 5E and Fig S5K). Importantly, both Pmel^shMettl3^ and Pmel^shDnmt3b^ T cells retained a significantly larger pool of pTex cells, marked by PD1⁺TCF1⁺ Vβ13⁺ expression (Fig 5F). Correspondingly, these cells demonstrated superior effector function, producing higher levels of IFN-γ and TNF-α upon in vitro re-stimulation (Fig S5L-S5M).

To directly evaluate anti-tumor efficacy in a therapeutic setting, three-day activated Pmel^Wt^, Pmel^shMettl3^ or Pmel^shDnmt3b^ cells were adoptively transferred into Rag1^⁻/⁻^ mice bearing established subcutaneous B16F10 tumors (Fig 5G). In this context as well, Pmel^shMettl3^ and Pmel^shDnmt3b^ cells conferred stronger tumor control than Pmel^Wt^ cells (Fig 5H-5I). Comprehensive profiling of Vβ13⁺ T cells revealed diminished frequencies of tTex subsets (PD1⁺Tim3⁺, PD1⁺CTLA4⁺, and PD1⁺CD39⁺) in Pmel^shMettl3^ and Pmel^shDnmt3b^ groups, compared to Pmel^Wt^ cells (Fig 5J-5K, Fig S5N-S5Q). Conversely, both Pmel^shMettl3^ and Pmel^shDnmt3b^ cells exhibited ∼4-fold enrichment of pTex cells (PD1⁺TCF1⁺), along with superior cytokine production (IFN-γ and TNF-α) compared with Pmel^Wt^ cells (Fig 5L-5O).

Collectively, these findings demonstrate that inhibiting Mettl3 or Dnmt3b generates long-lived, functionally competent CD8⁺ T cells that combine durable persistence with enhanced anti-tumor activity.

### Mettl3 or Dnmt3b knockdown renders Pmel T cells responsive to anti-PD1 therapy

The efficacy of anti-PD1 therapy depends on the exhaustion state of intratumoral T cells: tTex cells are largely refractory to treatment, whereas pTex cells can respond by proliferating and differentiating into effector subsets. Since the Mettl3-Dnmt3b axis restricts pTex differentiation, we next investigated whether targeting this pathway could enhance CD8⁺ T-cell responsiveness to anti-PD1 therapy.

Activated Pmel^Wt^, Pmel^shMettl3^, or Pmel^shDnmt3b^ cells were adoptively transferred into wild-type C57BL/6J mice bearing subcutaneously established B16F10 melanomas and treated with either anti-PD1 antibody or control IgG for the indicated time points (Fig 6A). Consistent with previous observations, tumor-reactive Pmel^Wt^ cells rapidly differentiated into tTex within the tumor and failed to respond to anti-PD1 therapy. In contrast, both Pmel^shMettl3^ and Pmel^shDnmt3b^cells mediated strong anti-tumor activity, which was further enhanced by anti-PD1 treatment (Fig 6B). Analysis of intratumoral Pmel T cells revealed that compared to Pmel^Wt^, the intratumoral frequencies of Pmel^shMettl3^ and Pmel^shDnmt3b^ cells were significantly higher, with modest additional increases upon PD1 blockade (Fig 6C). This improved persistence correlated with reduced differentiation into tTex (lower frequencies of PD1⁺Tim3⁺ and PD1⁺CD39⁺ subsets) (Fig 6D-6E), and maintenance of a pTex phenotype, as indicated by sustained TCF1 and FOXO1 expression (Fig 6F-6G). Functionally, Pmel^shMettl3^ and Pmel^shDnmt3b^ cells isolated from tumors also exhibited superior effector capacity, producing higher levels of IFN-γ and TNF-α upon re-stimulation compared to Pmel^Wt^ cells (Fig 6H-6I).

**Figure 6.**
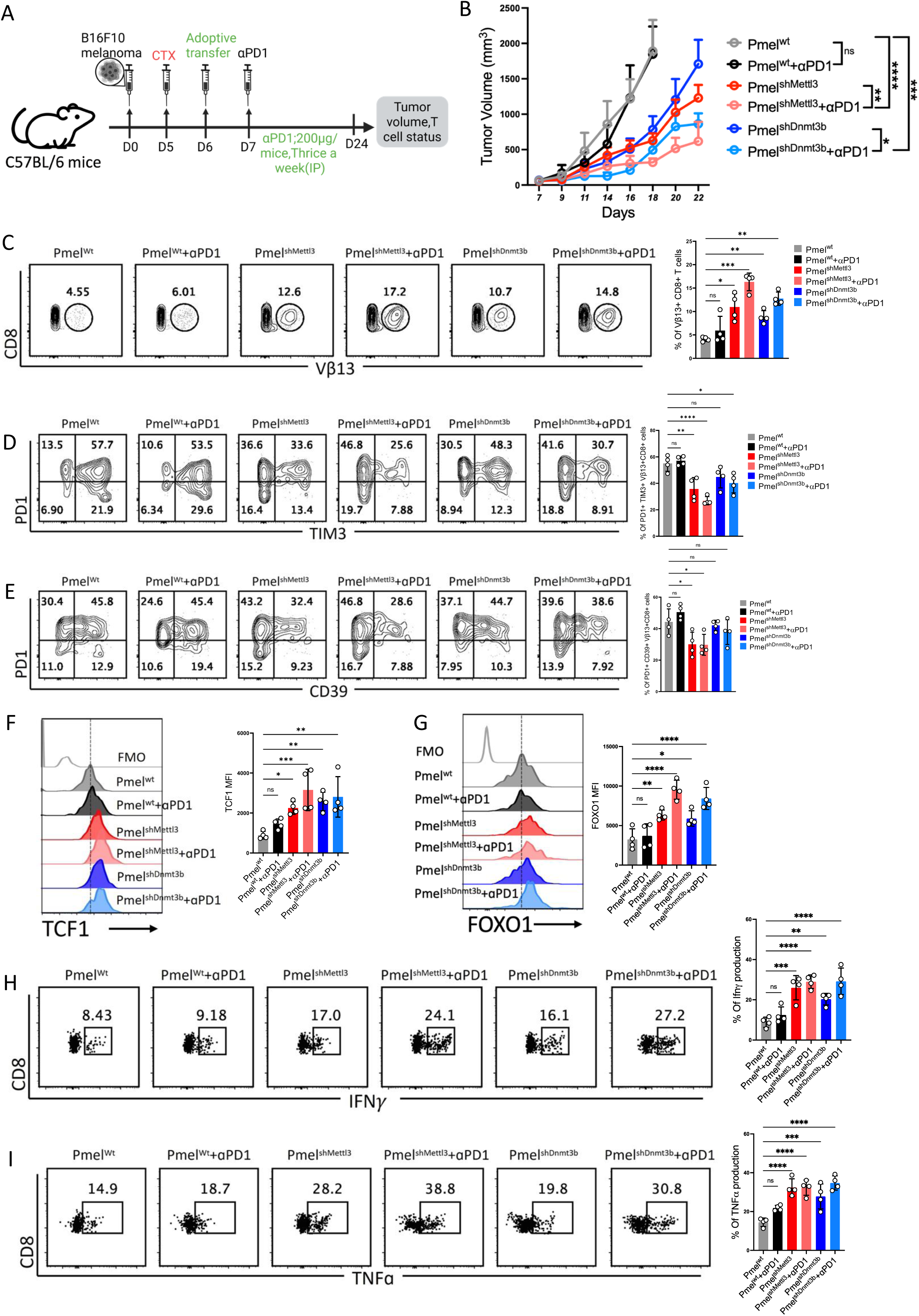
Genetic ablation of Mettl3 or Dnmt3b in CD8⁺ T cells enhances responsiveness to anti-PD1 therapy. (A) Schematic of the experimental design: C57BL/6J mice (N = 5 per group) were subcutaneously implanted with B16F10 melanoma and adoptively transferred with either wild-type Pmel (Pmel^Wt^), Mettl3-knockdown Pmel (Pmel^shMettl3^), or Dnmt3b-knockdown Pmel (Pmel^shDnmt3b^) T cells (1.5 × 10⁶ cells/mouse). Mice were subsequently treated with either control IgG or anti-PD1 antibody (200 μg/mouse, three times per week). (B) Tumor growth in mice from (A). (C-I) Adoptively transferred Pmel T cells (CD8⁺Vβ13⁺) isolated from tumors in (B) were analyzed for: (C) frequency; (D, E) cell surface expression of (D) PD1 and Tim3, and (E) PD1 and CD39; (F, G) intracellular expression of (F) TCF1, and (G) FOXO1; and (H, I) intracellular production of (H) IFNγ, and (I) TNFα, following in vitro re-stimulation. Adjacent bar graphs (C-I) represent cumulative data from N = 4 mice. *, P < 0.05; **, P < 0.01; ***, P < 0.005; ****, P < 0.0001.

Collectively, these results demonstrate that inhibiting Mettl3 or Dnmt3b preserves a pTex state, thereby sensitizing tumor-specific CD8⁺ T cells to anti-PD1 therapy and improving antitumor efficacy.

## Discussion

Although ICB, particularly anti-PD-1 and anti-CTLA-4, has transformed cancer immunotherapy by reinvigorating the functional capacity of Tex cells, the pronounced heterogeneity in clinical responsiveness underscores a pivotal and enduring question: are all Tex subsets equally amenable to therapeutic reprogramming? Recent studies have revealed a highly complex Tex landscape within the tumor microenvironment (TME), with subsets defined by distinct epigenetic and transcriptomic features and differing in their responsiveness to ICB(*6, 9*). While chronic antigen stimulation is the primary driver of Tex differentiation, it remains unclear which additional cellular cues guide the divergence into functionally distinct Tex lineages.

Here, we identify Mettl3 as a central epitranscriptomic regulator that dictates the balance between terminal exhaustion and memory-like differentiation in CD8⁺ T cells. We show that Mettl3 expression is enriched in tTex cells and inversely correlated with the pTex in both murine tumor models and chronically stimulated human CD8⁺ T cells. Mechanistically, Mettl3 promotes progressive exhaustion at the expense of memory formation by stabilizing *DNMT3B* transcripts through m⁶A modification. Elevated Dnmt3b in turn enforces CpG methylation and chromatin compaction at memory-associated loci, transcriptionally silencing programs required for sustaining progenitor and multipotent states.

Our data reveal an intriguing observation that the critical decision between terminal differentiation and memory precursor generation occurs early during CD8⁺ T cell activation and is largely governed by Mettl3 expression. This is further exemplified by the skewed differentiation of TCF1⁺ self-renewing progenitor cells under Mettl3-attenuated conditions, even when CD8⁺ T cells were chronically stimulated and strongly driven toward the formation of terminal Tex (tTex) cells. Recent studies mapping the Tex differentiation trajectory in chronic viral infection models highlight the early emergence of pTex cells with distinct epigenetic and transcriptional features(*8, 38*). These pTex cells are proposed to seed tTex cells through an intermediate Tex (Tex^int^) population marked by CX3CR1 expression as exhaustion progresses(*8, 39*). While this model supports a linear differentiation pathway from pTex to Tex^int^ and subsequently to tTex, other reports suggest a divergent route, showing that pTex, although established early, retain ‘transcriptional similarity’ even at later stages of chronic infection(*38*). Within this context, our data provide mechanistic insights into the development of progenitor populations and their differentiation into progeny subsets. Based on our findings, we envisage that during the initial TCR-mediated activation phase, the relative abundance of Mettl3 shapes the early heterogeneity between effector and progenitor CD8⁺ T cell subsets. Subsequently, under chronic antigenic stimulation, as seen in tumors or persistent viral infections, a fraction of progenitor cells may preserve their identity by maintaining low Mettl3 levels, whereas others progressively acquire Mettl3 and diversify into distinct Tex subsets. Although the environmental and cellular cues that regulate Mettl3 expression in CD8⁺ T cells remain to be elucidated, our findings suggest that Mettl3 may function as a molecular rheostat, fine-tuning the balance between progenitor maintenance and progeny differentiation.

Diving deeper, we found that Mettl3-mediated progenitor bifurcation is primarily driven by chromatin remodeling at memory-associated gene loci, rather than by altered accessibility at exhaustion- or effector-associated loci. This indicates that Mettl3 inhibition may not yield a uniform progenitor CD8⁺ T cell pool but instead generates a spectrum of progenitors epigenetically primed for distinct phenotypic and functional outcomes. Future studies integrating scRNA-seq with scATAC-seq will be instrumental in resolving these developmental trajectories with higher precision. Mechanistically, we show that Mettl3-dependent chromatin remodeling in CD8⁺ T cells is largely mediated by Dnmt3b, which enforces CpG methylation, chromatin compaction, and transcriptional repression. We further identify Dnmt3b as the principal downstream effector of Mettl3: by stabilizing *DNMT3B* transcripts via m⁶A modification, Mettl3 sustains DNA methylation and closed chromatin states. Previous studies have also implicated Mettl3 in regulating DNA methylation and chromatin accessibility across diverse cell types(*40-43*). However, whereas those reports emphasized the role of Mettl3 in recruiting Dnmt1 or the DNA demethylase TET1 to shape the 5mC landscape(*42, 43*), our findings uncover an additional mechanism-m⁶A-dependent stabilization of *DNMT3B* transcripts. Notably, de novo DNA methylation by Dnmt3a has recently been implicated in reinforcing terminal exhaustion features(*44*). Although we did not observe m⁶A-dependent regulation of *DNMT3A* stability, it remains important to investigate whether Mettl3 plays an ancillary role in modulating Dnmt3a activity and promoter recruitment in Tex cells.

Building on our finding that the Mettl3-m⁶A-Dnmt3b axis restricts progenitor CD8⁺ T cell formation, we performed a “T cell parking” experiment in Rag1^⁻/⁻^ mice to directly test whether loss of Mettl3 or Dnmt3b preserves memory potential. Knockdown of either gene not only enhanced homeostatic proliferation but also enabled robust recall responses and effective control of B16F10 melanoma, demonstrating their capacity to rapidly transition into effector cells, a defining feature of memory T cells(*45*). Under tumor rechallenge, a significant fraction of Pmel T cells transduced with Mettl3 or Dnmt3b shRNA maintained a PD1⁺TCF1⁺ pTex phenotype while simultaneously generating a more differentiated PD1⁺TCF1⁻ subset, consistent with chronic infection models where progenitor-like T cells ensure long-term persistence and sustained effector output(*38*). The self-renewing potential conferred by inhibition of the Mettl3-m⁶A-Dnmt3b pathway was further supported by superior responsiveness to anti-PD1 therapy, underscoring the translational value of targeting this axis to improve the depth and durability of ICB responses in human cancers. Early-phase clinical trials should focus on evaluating selective Mettl3 or Dnmt3b inhibitors, alone or in combination with PD-1 blockade, in patients with tumors characterized by profound T cell exhaustion. Stratification of patients based on Tex subset composition and Mettl3 expression levels may further guide precision therapy. Ultimately, integrating epitranscriptomic modulators with existing immunotherapies has the potential to overcome resistance, sustain long-term antitumor immunity, and significantly broaden the clinical benefit of ICB. Notably, a recent study reported that pharmacological inhibition of Mettl3 improves anti-PD1 responses via distinct mechanisms, including induction of type I interferons and increased tumor immunogenicity(*46*). Our results extend these observations by establishing a CD8⁺ T cell-intrinsic role of Mettl3 in restricting progenitor potential, thereby providing a mechanistic rationale for targeting this pathway to improve immune checkpoint blockade outcomes.

In summary, we identify Mettl3 as a molecular rheostat of CD8⁺ T cell fate, where its m6A-dependent stabilization of Dnmt3b drives terminal exhaustion at the expense of progenitor maintenance. Disruption of this axis preserves self-renewing potential, enhances recall responses, and improves sensitivity to PD-1 blockade, establishing epitranscriptomic control of T cell differentiation as a tractable target to sustain durable antitumor immunity.

## Materials and Methods

### Mice

C57BL/6J, Pmel and Rag1⁻/⁻ mice were obtained from Jackson Laboratory (Bar Harbor, MA) and maintained under pathogen-free conditions at CSIR-IICB. All animal experiments were conducted in accordance with institutional guidelines and approved by the Institutional Animal Ethics Committee, CSIR-IICB, Kolkata, India. For tumor studies, 6-8-week-old male and female mice were randomly assigned to experimental groups, and no sex-related differences in outcomes were observed.

### Cell Lines

B16-F10, HEK293T, YUMM1.7, and MC38 cell lines were obtained from ATCC and confirmed Mycoplasma-free using the MycoAlert Kit (Lonza). Cells between passages 4-6 were used for mouse injections and lentiviral production.

### In vitro activation and culture of human CD8+ T cell

De-identified buffy coats were used in this study under Institutional IRB approval (IICB/IRB/2020/2P). PBMCs were isolated from healthy donor buffy coats by Ficoll-Hypaque density-gradient centrifugation, and CD8⁺ T cells were purified by negative selection using Dynabeads (Invitrogen, USA). Cells were activated for 72 h with plate-bound anti-CD3 (5 µg/mL) and anti-CD28 (2 µg/mL) in RPMI-1640 supplemented with 10% FBS, penicillin-streptomycin, and IL-2 (100 U/mL). For Tex differentiation, activated cells were cultured for 12 days under chronic TCR stimulation (plate-bound anti-CD3, IL-2 at 25 U/mL), with passaging every 48 h. For acute stimulation, activated CD8⁺ T cells were cultured for 12 days with IL-2 (100 U/mL) but without TCR stimulation. For memory T cell (T_Mem_) differentiation, activated cells were maintained for 12 days in RPMI-1640/10% FBS supplemented with IL-15 (10 ng/mL) and IL-7 (10 ng/mL), also without TCR stimulation. In some experiments, CD8⁺ T cells were activated in the presence or absence of STM2457 (10 µM) or nanaomycin A (DNMT3Bi; 250 nM) before being differentiated into either Tex or TMem. All groups were restimulated overnight with plate-bound anti-CD3 (5 µg/mL) prior to analysis.

### Adoptive T Cell Transfer

Wild type (Wt) or Rag1^-/-^ C57BL/6J (B6) mice of 6-8 weeks of age were subcutaneously implanted with B16-F10 melanoma cells (0.5 × 10⁶). Eight days later, mice were lymphodepleted (in case of Wt B6 mice) with cyclophosphamide (4 mg/mouse) and subsequently received adoptive transfer (i.v.) of Pmel T cells (1.5 × 10⁶/mouse) that had been activated for two days with gp100 peptide (3 μg/mL) and transduced with control shRNA, shMettl3, or shDnmt3b. IL-2 (50,000 U/mouse; i.p.) was administered for three consecutive days, and in some experiments, mice were additionally treated with anti-PD1 antibody (200 μg/mouse) or control IgG three times weekly.

### Tumor-infiltrating T cell isolation

Tumors from B16-F10, MC38, and YUMM1.7-bearing mice were aseptically excised and transferred to cold RPMI-1640. Tissues were minced with sterile scalpels and digested in RPMI-1640 containing collagenase type IV (2 mg/mL) and DNase I (100 µg/mL) at 37 °C for 45 min with gentle agitation. Digests were passed through a 70 µm strainer to generate single-cell suspensions, followed by ACK lysis if required. TILs were enriched by density centrifugation over Hi-Sep LSM at 1200 rpm for 30 min at room temperature. Mononuclear cells at the interface were collected, washed, and resuspended in RPMI-1640 for downstream use.

### Flow cytometry staining and analysis

Cells were incubated with fluorochrome-conjugated antibodies in FACS buffer (0.5% BSA, 0.1% sodium azide in PBS) for 30 min at 4 °C, washed, and resuspended in FACS buffer with 1% paraformaldehyde for staining surface molecules. For intracellular cytokine staining (IFN-γ, TNF-α), T cells were restimulated for 4 h at 37 °C with PMA (100 ng/mL), ionomycin (1 µg/mL), and Golgi Plug (BD Biosciences), followed by surface staining, fixation/permeabilization (BD Cytofix/Cytoperm), and intracellular staining. For nuclear proteins, surface staining was followed by fixation/permeabilization using the FoxP3 Staining Buffer Set (Thermo Fisher Scientific). A fixable viability dye (Live/Dead Fixable Yellow, Thermo Fisher Scientific) was used after surface marker staining. Samples were acquired on a BD LSR Fortessa II (BD Biosciences) and analyzed with FlowJo software.

### Extracellular metabolic flux assay

T cell OCR and ECAR were measured using the Agilent Seahorse XFe24 Analyzer. Activated T cells (3 × 10⁵/well) were seeded in Cell-Tak–coated plates in XF Base Medium (1 mM pyruvate, 1% FBS; ±10 mM glucose for OCR/ECAR, pH 7.4), centrifuged (200 × g, 1 min), and incubated 30 min at 37°C in a non-CO₂ incubator. OCR and ECAR were assessed following sequential injections: oligomycin (10 μM), FCCP (10 μM), rotenone/antimycin A (1 μM each) for OCR; glucose (10 mM), oligomycin (10 μM), 2-DG (100 mM) for ECAR, using standard Seahorse cycles (mix 3 min, wait 2 min, measure 3 min). Data were analyzed with Wave software (Agilent).

### Western blot analysis

T cells were lysed in RIPA buffer with protease and phosphatase inhibitors (Takara), incubated on ice 30 min, and cleared by centrifugation (14,000 × g, 15 min, 4°C). Protein concentration was measured by Bradford assay, mixed with Laemmli buffer, boiled (95°C, 5 min), and resolved by SDS-PAGE. Proteins were transferred to PVDF membranes, blocked in 5% BSA/TBST (1 h, RT), and incubated overnight at 4°C with primary antibodies (1:1000), followed by HRP-conjugated secondary antibodies (1:2000, 1 h, RT). Bands were detected by ECL (Bio-Rad) and imaged using a ChemiDoc system; quantification was performed with ImageJ.

### Prediction of m^6^A sites in *DNMT3B* mRNA

The intron-excluded human DNMT3B mRNA (FASTA, NCBI) was analyzed with SRAMP, which integrates three Random Forest classifiers—positional nucleotide patterns, KNN, and nucleotide pair spectrum features—to assign site-specific confidence scores. Fourteen putative m⁶A sites were identified: high (H1–H2), moderate (M1–M5), and low (L1–L7) confidence. Primers flanking each site were designed via Primer-BLAST, and an internal region lacking predicted m⁶A sites served as a negative control (m⁶A-neg); sequences and scores are listed in Table 1.

**Table1:**
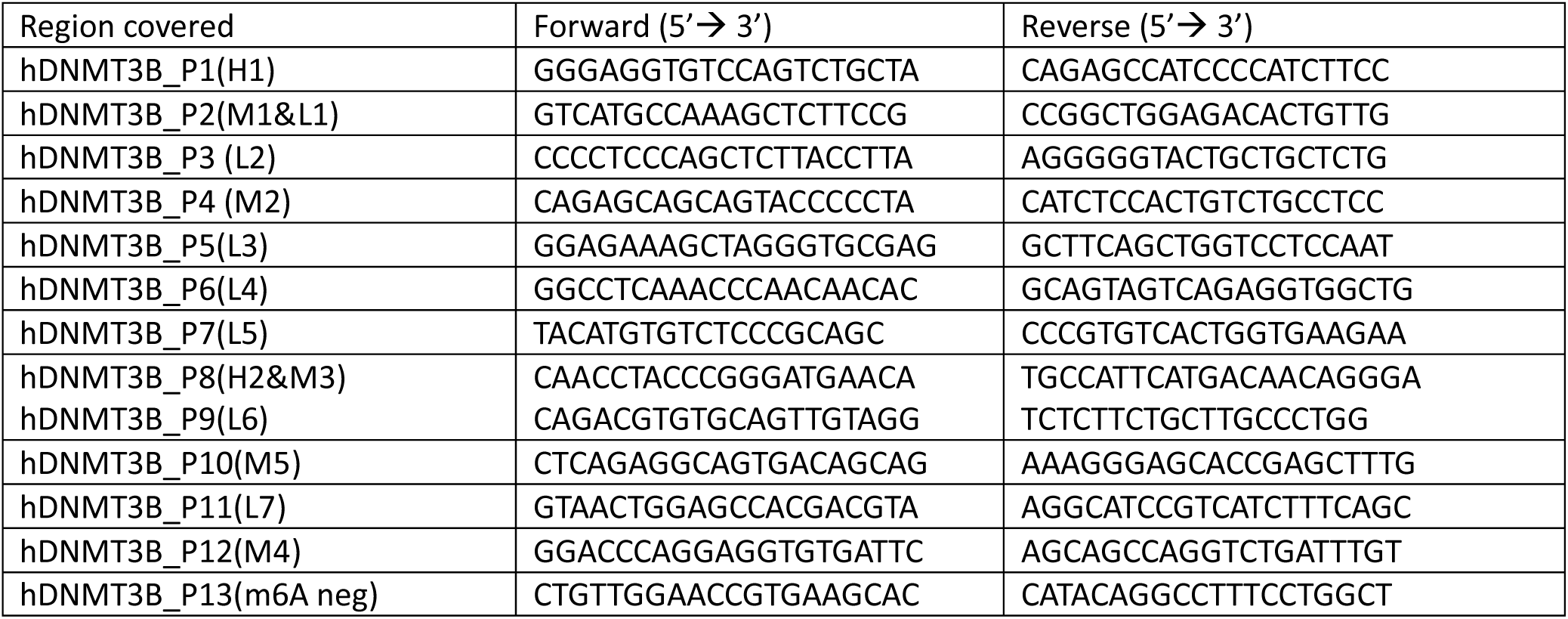
RNA-IP primers.

### m^6^A RNA Immunoprecipitation

m⁶A RIP was performed using the Magna MeRIP™ m⁶A Kit (Millipore, Cat. 17-10499) per manufacturer’s instructions. Total RNA from human CD8⁺ T cells was extracted using RNAqueous Phenol-free kit (Invitrogen, Cat. AM1912). For each reaction, 300 µg RNA was fragmented (∼100 nt) in 10× Fragmentation Buffer at 94 °C for 4 min and quenched with 0.5 M EDTA; fragment size was verified on 1.5% agarose gel. Protein A/G magnetic beads were pre-washed in 1× IP Buffer and incubated 30 min at RT with 10 µg anti-m⁶A antibody or mouse IgG. Fragmented RNA was incubated with antibody-bound beads 2h at 4 °C with rotation, washed thrice with cold IP Buffer, and m⁶A-enriched RNA eluted with 20 mM N⁶-methyladenosine 5′-monophosphate in Elution Buffer and purified. Ten percent of fragmented RNA was retained as input for normalization. Enriched RNA and inputs were analyzed by RT-qPCR to assess target region amplification.

### CpG island prediction,primer design and bisulfite conversion

A 4 kb upstream genomic sequence from transcriptional start site (TSS) of the target gene was retrieved from NCBI for CpG island analysis using MethPrimer. CpG islands were predicted using criteria of >100 bp, GC >50%, and observed/expected CpG ratio >0.6, and MSPs were designed ensuring ≥6 CpG sites per region. Genomic DNA from human CD8⁺ T cells was bisulfite-converted using the EZ DNA Methylation-Gold Kit (Zymo Research) per manufacturer’s instructions: 300 ng DNA was treated to convert unmethylated cytosines to uracils, followed by denaturation, bisulfite conversion, desulphonation, and purification; converted DNA was eluted in 20 µL and stored at −20 °C. Designed methylation and unmethylation-specific primers for PCR analysis are listed in Tables 2 and 3.

**Table 2:**
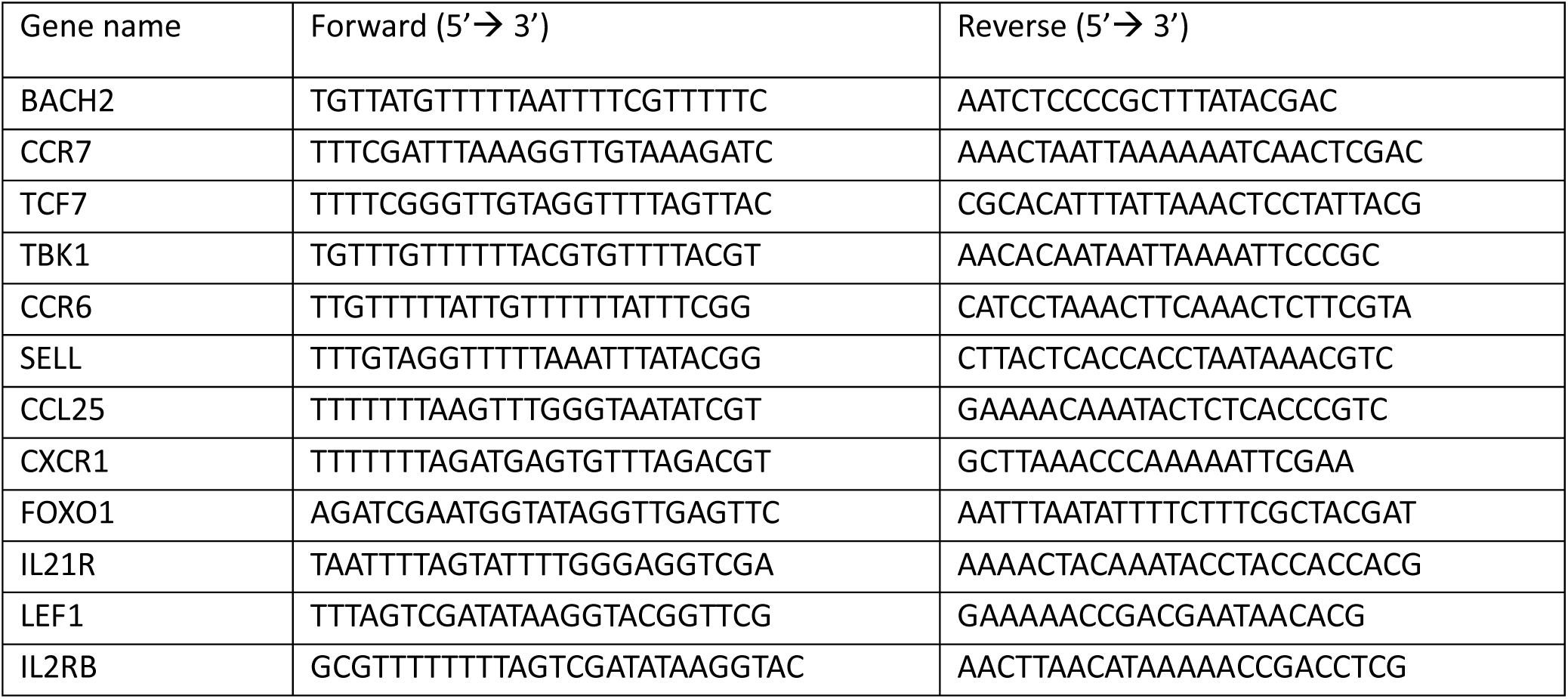
Methylation specific primers.

**Table 3:**
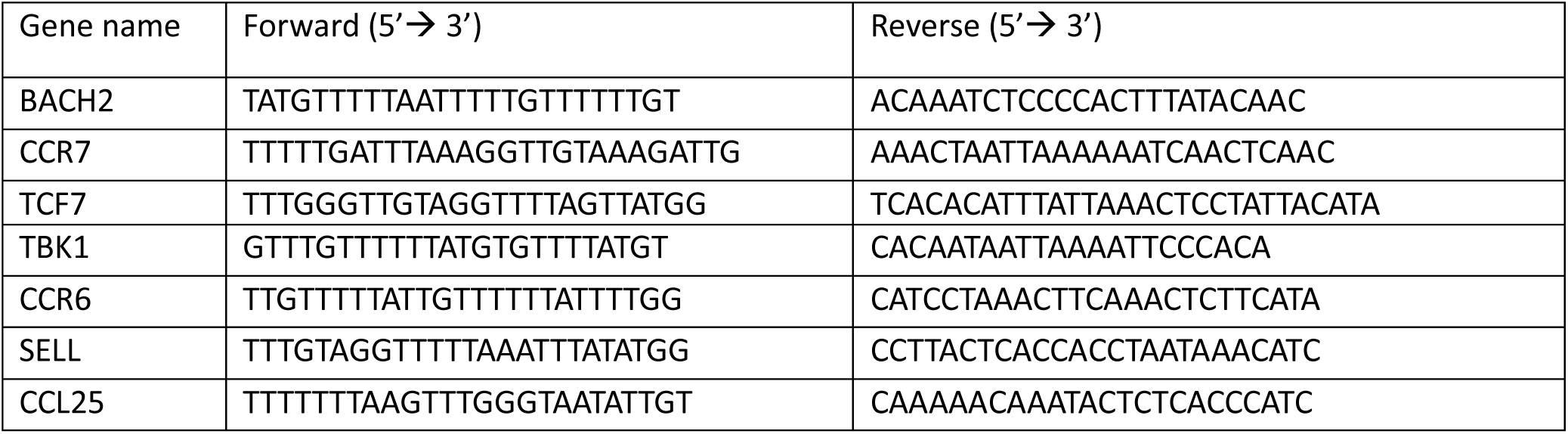
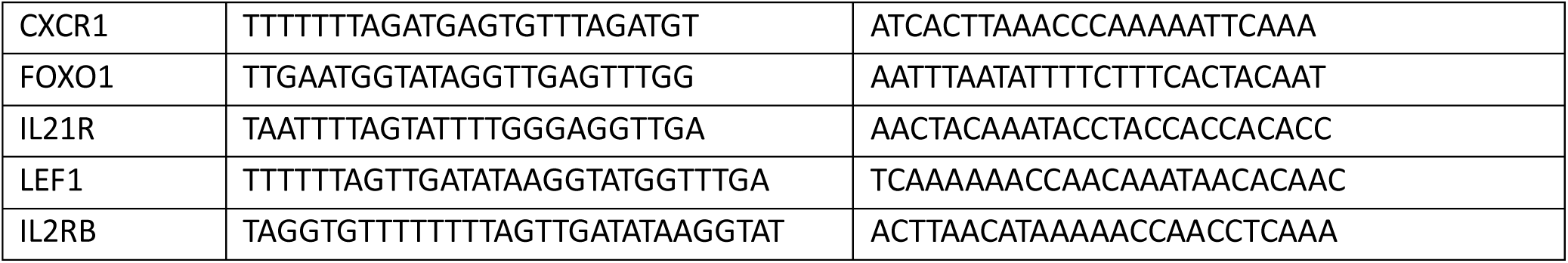
Unmethylation specific primers.

### Quantification of m^6^A RNA methylation

Global N6-methyladenosine (m6A) levels were quantified using the EpiQuik™ m6A RNA Methylation Quantification Kit (Colorimetric; Epigentek), following the manufacturer’s protocol. Briefly, 200 ng of total RNA isolated from human CD8⁺ T cells was added to strip wells pre-coated with binding solution and incubated at 37 °C to ensure efficient immobilization. After washing, wells were sequentially treated with an m6A-specific capture antibody, detection antibody, and enhancer solution, with wash steps performed between each incubation to minimize background. Colorimetric signal was developed by adding the developer solution, and the reaction was stopped with stop solution once adequate color formation was observed. Absorbance was measured immediately at 450 nm using a microplate reader. Negative and positive controls (m6A RNA standard) supplied with the kit were included in each assay for normalization. The relative percentage of m6A in RNA samples was calculated using the formula: 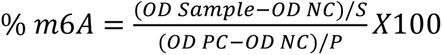 where *OD* is the optical density at 450 nm, *NC* is the negative control, *PC* is the positive control, *S* is the amount of input sample RNA (ng), and *P* is the amount of positive control RNA (ng). In experiments where a standard curve was generated, absolute m6A content (ng) was derived from the linear regression of OD values and expressed as a percentage of total RNA.

### Lentiviral transduction of CD8+ T cell

HEK293T cells (ATCC) were cultured in T75 flasks at 37 °C, 5% CO₂ in DMEM (Gibco) supplemented with 10% FBS and 1% penicillin–streptomycin. For virus production, cells were co-transfected with 15 µg of lentiviral expression plasmids encoding METTL3 shRNA, METTL3 overexpression, DNMT3B shRNA, or non-targeting control (Origene), together with 10 µg psPAX2 and 5 µg pMD2.G, using CaCl₂/HBS precipitation. After 24 h, medium was replaced with low-serum medium, and supernatant was collected at 48–72 h, clarified (500 × g, 5 min), filtered (0.45 µm, Millipore), concentrated using Lenti-X (Takara Bio), aliquoted, and stored at −80 °C. Activated CD8⁺ T cells were washed, plated on RetroNectin-coated 24-well plates (30 µg/mL, 4 °C overnight), and transduced by spinoculation (1,000 × g, 1.5 h, 32 °C) in the presence of IL-2 (100 U/mL), followed by 24 h culture at 37 °C, 5% CO₂.

### RNA extraction, cDNA synthesis, qPCR

Total RNA was extracted with TRIzol (Invitrogen), purified, and resuspended in nuclease-free water. RNA concentration and purity were measured using a Multiskan SkyHigh spectrophotometer. cDNA (1 µg RNA) was synthesized using iScript (Bio-Rad) with random primers and oligo(dT), diluted 1:3, and amplified by qPCR using iTaq Universal SYBR Green (Bio-Rad) on a CFX96 system. Gene-specific primers were designed via NCBI Primer-BLAST (Table X). Relative expression was calculated using ΔΔCt with 18S rRNA or β-actin as housekeeping controls.

### mRNA stability assay

Activated CD8⁺ T cells were treated with or without the METTL3 inhibitor STM2457, followed by transcriptional arrest with Actinomycin D (5 µg/mL). Cells were harvested at 0, 2, 4, 6, 8, and 10 h; RNA was extracted and reverse transcribed as above. qPCR was performed using β-actin as control. Decay kinetics were modelled in GraphPad Prism v9 using one-phase exponential decay, non-linear regression (least squares, 95% CI, asymmetrical confidence intervals), and medium convergence. Decay rate and half-life were derived from fitted curves.

### Cell cycle analysis

CD8⁺ T cells were pulsed with BrdU (10 µM) overnight, fixed/permeabilized (BD buffer), and treated with DNase I (Sigma-Aldrich, in DPBS with Ca²⁺/Mg²⁺, 37 °C, 1 h). BrdU incorporation was detected with APC-conjugated anti-BrdU antibody; total DNA was stained with 7-AAD in RNase A. Samples were acquired on a BD LSRFortessa™ and analyzed with FlowJo to quantify G0/G1, S, and G2/M phases.

### RT^2^ profiler PCR array

Total RNA from human CD8⁺ T cells activated ± STM was isolated using the RNAqueous Phenol-free Kit. RNA purity and concentration were assessed spectrophotometrically (A₂₆₀/A₂₈₀ ∼1.8– 2.0; A₂₆₀/A₂₃₀ >1.7). For each sample, 0.5 µg RNA was reverse-transcribed with the RT² First Strand Kit (QIAGEN) including genomic DNA elimination (42 °C, 5 min), reverse transcription (42 °C, 15 min), and enzyme inactivation (95 °C, 5 min). cDNA was diluted and profiled using the RT² Profiler™ PCR Array Human Epigenetic Chromatin Modification Enzymes (PAHS-085ZD-6) with RT² SYBR® Green qPCR Mastermix under recommended cycling. Arrays included five housekeeping genes, three reverse-transcription, three positive PCR, and one genomic DNA control; accepted if genomic DNA Cₜ ≥ 35, positive PCR Cₜ = 20 ± 2, and reverse-transcription controls were consistent. Data were normalized to housekeeping genes and analyzed via ΔΔCₜ using QIAGEN tools.

### RNA extraction, library preparation, and sequencing

Human CD8⁺ T cells were activated ±STM, washed with PBS, and lysed in TRIzol. Total RNA was extracted using the RNeasy Mini Kit (Qiagen, Cat# 74104) and quality assessed by Qubit, NanoDrop, and TapeStation. Eight high-quality samples (Control and STM; n=4/group from two donors) were used for library preparation with the KAPA RNA HyperPrep Kit (Roche, Cat# 0000141759). Poly(A)⁺ RNA was enriched with oligo(dT) beads, fragmented (94 °C, 6 min), and converted to strand-specific cDNA. After A-tailing, adapter ligation, and purification, libraries were PCR-amplified (13 cycles), purified, and checked for fragment size using the Agilent dsDNA Reagent Kit (1–6000 bp) before sequencing. Sequencing was performed on a NovaSeq S4 sequencer with a read length of 2×150 bp and a target of 40 million reads per sample.

### Bulk RNA-Seq analysis

Raw RNA-seq reads were first subjected to quality assessment and adapter trimming using Trim Galore to remove low-quality bases and sequencing artifacts. The cleaned reads were then aligned to the human reference genome (GRCh38) using the STAR aligner(*47*), which allows rapid and accurate spliced-read mapping. Gene-level counts were quantified with featureCounts(*48*) to produce a raw count matrix. The count matrix, along with sample metadata, was subsequently imported into R for downstream analysis. Differential gene expression was evaluated with the DESeq2 package(*49*), which applies shrinkage estimators for dispersion and fold-change to improve statistical power and control false discovery rates. Genes with an adjusted *P* value (FDR) < 0.05 were considered significantly differentially expressed. Data visualization was performed in R, including principal component analysis to assess sample clustering and pheatmap for generating expression heatmaps of differentially expressed genes.

### Single-cell RNA-seq data analysis

Single-cell RNA-seq data were analyzed using Seurat (v4.3.0) in R(*50*). Gene expression matrices from each sample were imported, and low-quality cells were excluded based on mitochondrial transcript content and detected feature thresholds. Data were normalized with the LogNormalize method, and highly variable genes were identified using *FindVariableFeatures* (“vst”). The datasets were merged, scaled (*ScaleData*), and subjected to principal component analysis (PCA), followed by batch-effect correction with Harmony (v1.0)(*51*). Cells were clustered using a shared nearest-neighbor graph with the Louvain algorithm and visualized in two dimensions using t-SNE and UMAP. Cluster-specific marker genes were identified with *FindAllMarkers*, and the top 10 differentially expressed genes per cluster were extracted to define transcriptional identities. Module scoring (*AddModuleScore*) was applied to assess pathway or gene set activity across clusters, and expression patterns were visualized with feature plots and dot plots to capture transcriptional heterogeneity.

### ATAC-seq sample preparation and sequencing

Human peripheral blood CD8⁺ T cells were isolated from healthy donors and cultured under activation conditions for 3 days in the presence or absence of STM. Cells were divided into two groups (control and STM-treated), each consisting of three biological replicates. Nuclei were isolated from 5 × 10⁵ cells per sample using the ATAC-Seq Kit (Active Motif, Cat. No. 53150) according to the manufacturer’s instructions. The isolated nuclei underwent tagmentation with Tn5 transposase, DNA purification, and library amplification. A total of six ATAC-seq libraries were generated and sequenced on an Illumina NovaSeq platform with paired-end 2 × 100 bp reads, yielding approximately 50 million reads per sample.

### ATAC-seq data analysis

ATAC-seq reads were first quality-checked, trimmed for adapters and low-quality bases using Trim Galore, and aligned to the human reference genome (GRCh38) with Bowtie2(*52*). Aligned reads were processed with Samtools(*53*) to remove duplicates and mitochondrial reads, followed by filtering for properly paired, high-quality fragments. Picard was used to mark duplicates, and blacklist regions (ENCODE hg38) were removed with Bedtools(*54*). Peaks of accessible chromatin were called using MACS2(*55*) in paired-end mode. Peak sets were merged across samples to obtain consensus regions, and read counts were assigned to peaks using featureCounts(*48*). Peaks were annotated to nearby genes with HOMER(*56*), and downstream analyses—including differential accessibility, visualization etc-were carried out in R.

## Data availability

Mouse single-cell RNA-sequencing data were obtained from NCBI GEO (GSM6704035: CD8⁺ T cells from B16 melanoma tumor in wild-type mice; GSM6704043: CD8⁺ T cells from tumor-draining lymph node in wild-type mice). All human single-cell RNA-sequencing data analysed in this study were obtained from the 10x Genomics public repository We used datasets from 320k scFFPE from 8 Human Tissues (https://www.10xgenomics.com/datasets/320k_scFFPE_16-plex_GEM-X_FLEX) and 40k Mixture of Dissociated Tumor Cells from 4 Donors (https://www.10xgenomics.com/datasets/40k-mixture-of-dissociated-tumor-cells-from-4-donors-multiplexed-samples-4-probe-barcodes-1-standard), which include Skin Melanoma (BC13–14), Breast Cancer (BC2), Colorectal Cancer (BC3–4), and Endocervical Adenocarcinoma (BC15–16). These datasets are publicly available and fully described by respective repositories.

## Statistical Analysis

All experiments were performed with a minimum of three independent biological replicates. Data are presented as mean ± standard error of the mean (SEM) unless otherwise stated. Statistical comparisons between two groups were performed using unpaired two-tailed Student’s t-test, while multiple group comparisons were analyzed using one-way or two-way ANOVA followed by appropriate post hoc tests. A p-value < 0.05 was considered statistically significant. Graphs and statistical analyses were generated using GraphPad Prism or R.

## Author contributions

Conceptualization: PG and SC (Shilpak Chatterjee); Methodology: PG, AG, DB, SM, SC (Soham Chowdhury), IS, AM, AK, SC (Snehanshu Chowdhury) SP, and SC (Shilpak Chatterjee); Investigation: PG, DB, SM, SC (Soham Chowdhury), and IS; Visualization: PG, AG, SP and SC (Shilpak Chatterjee); Formal analysis: PG, AG, SP, and SC (Shilpak Chatterjee); Software: PG, AG, and SP; Funding acquisition: SC (Shilpak Chatterjee); Project administration: SC (Shilpak Chatterjee); Supervision: SC (Shilpak Chatterjee); Writing-original draft: SC (Shilpak Chatterjee).

## Tables

**Table 4:**
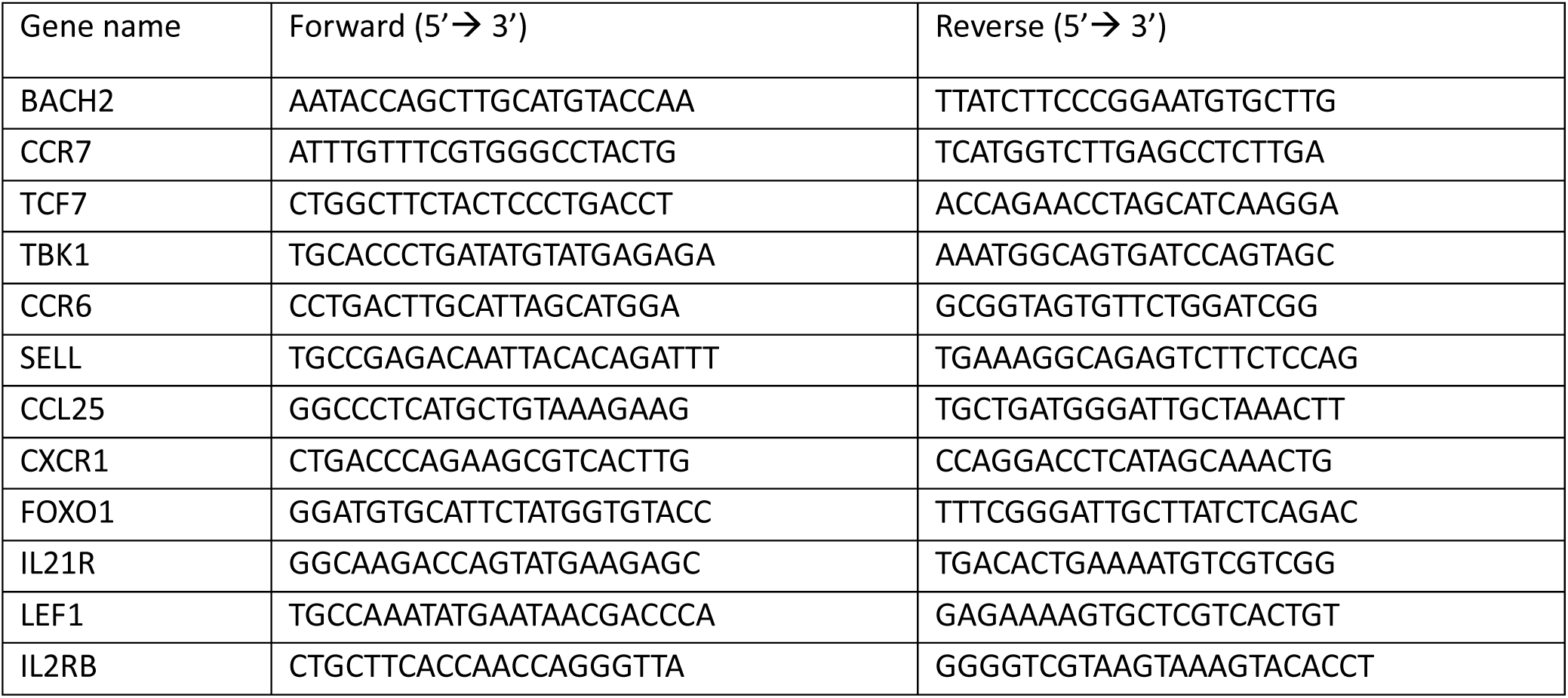
Gene specific primers for qPCR.

**Table 5:**
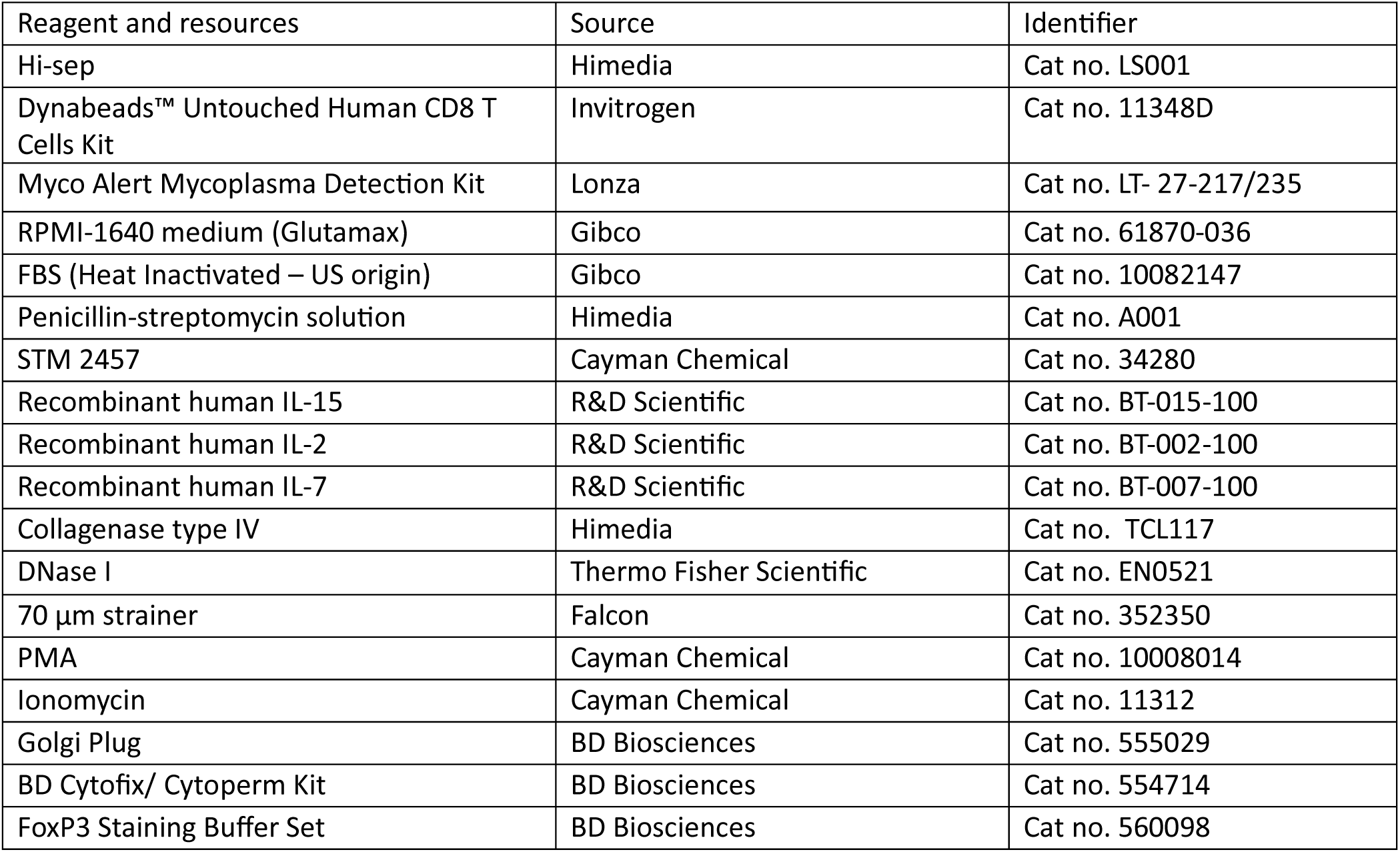
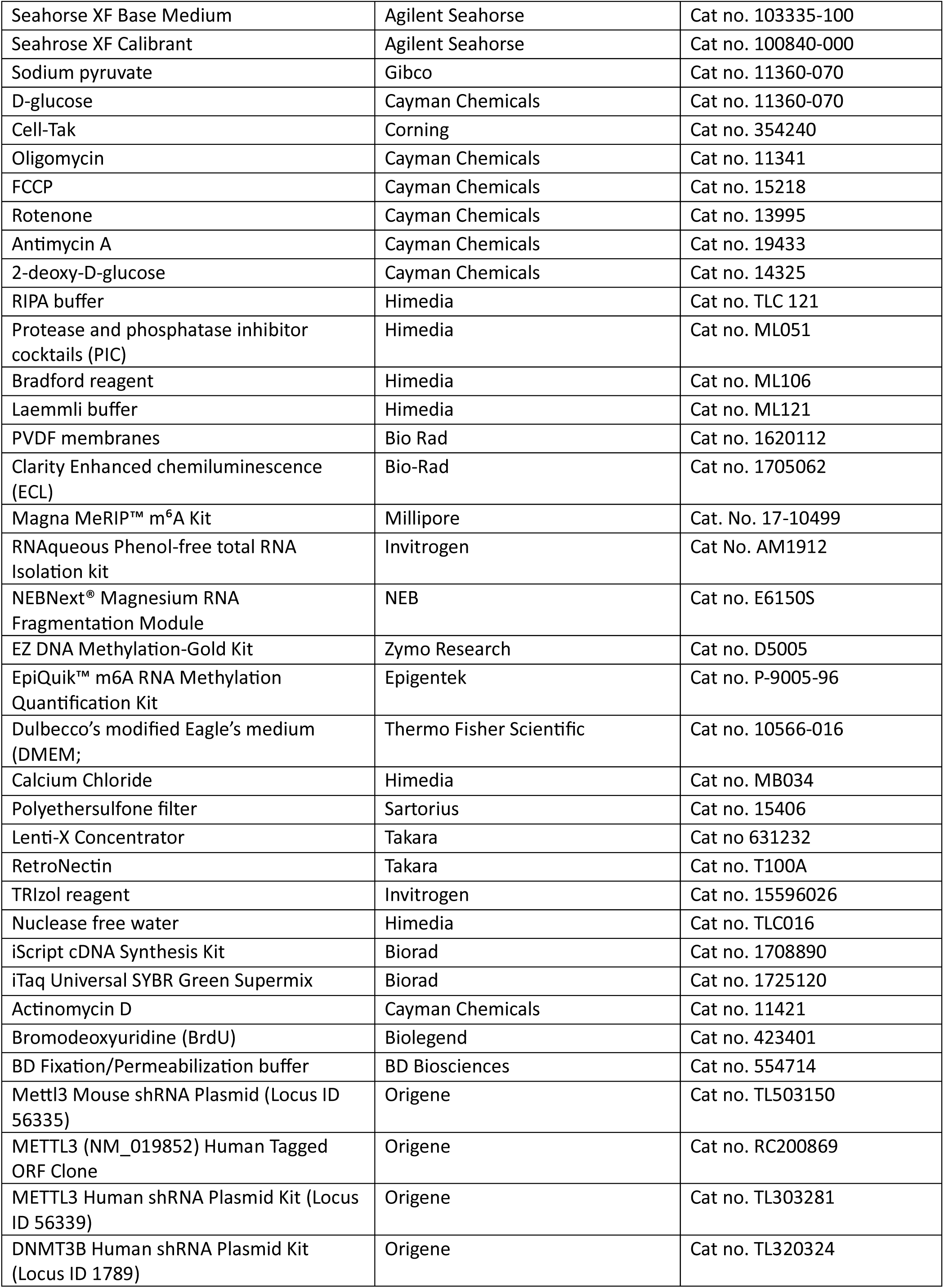
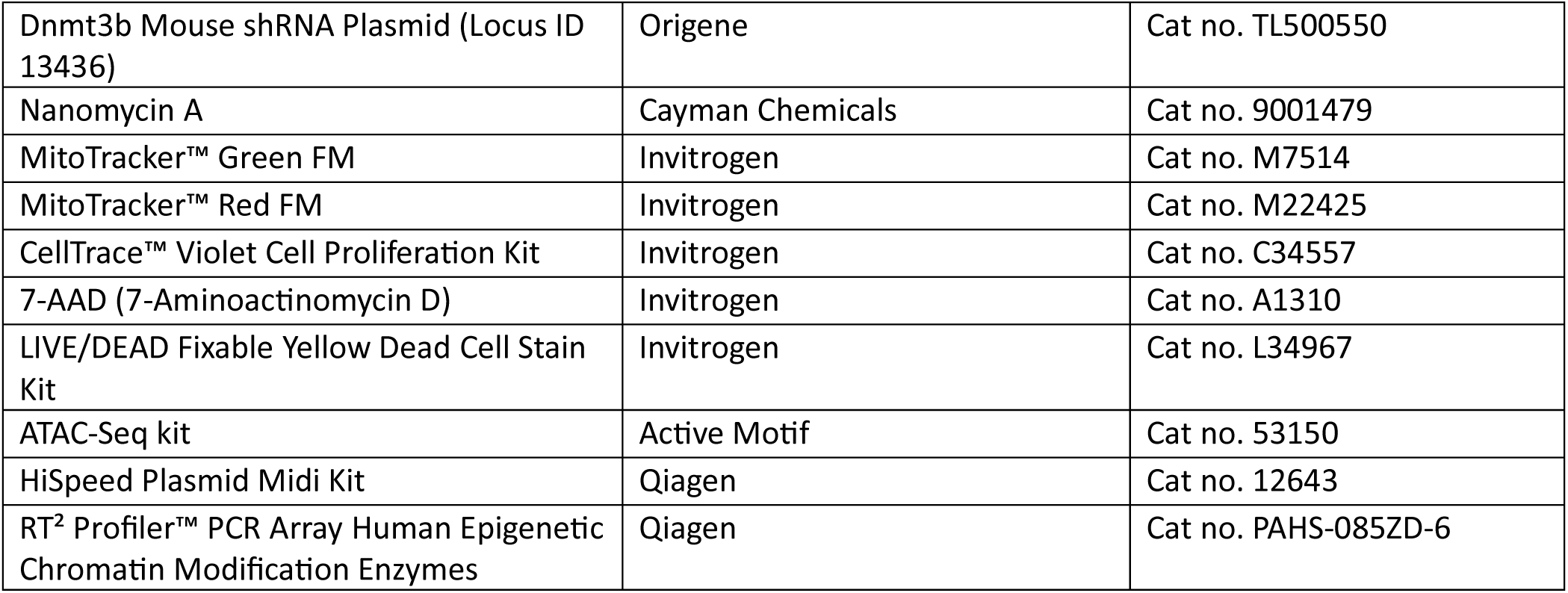
Reagent and resources.

**Table 6:**
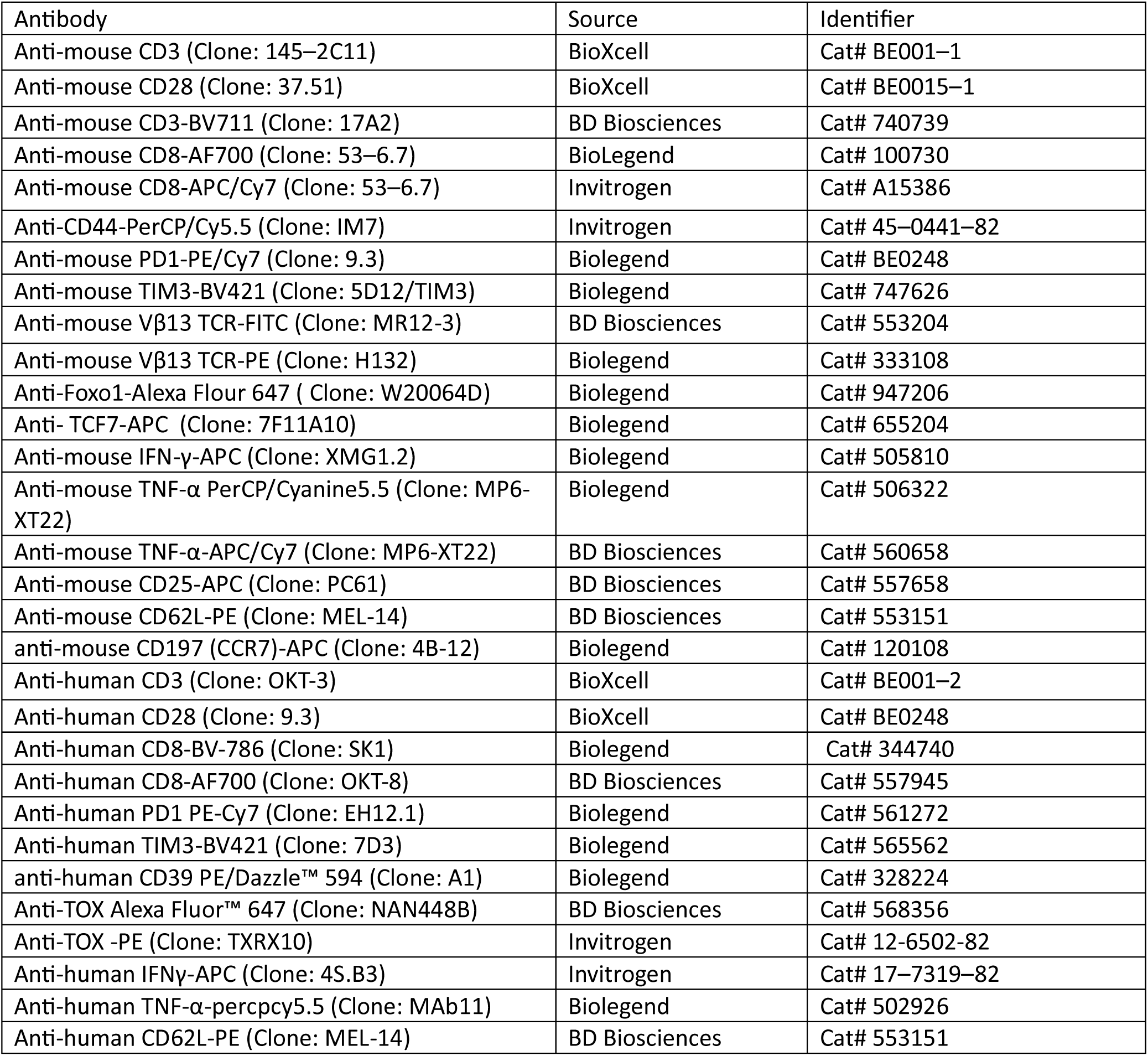
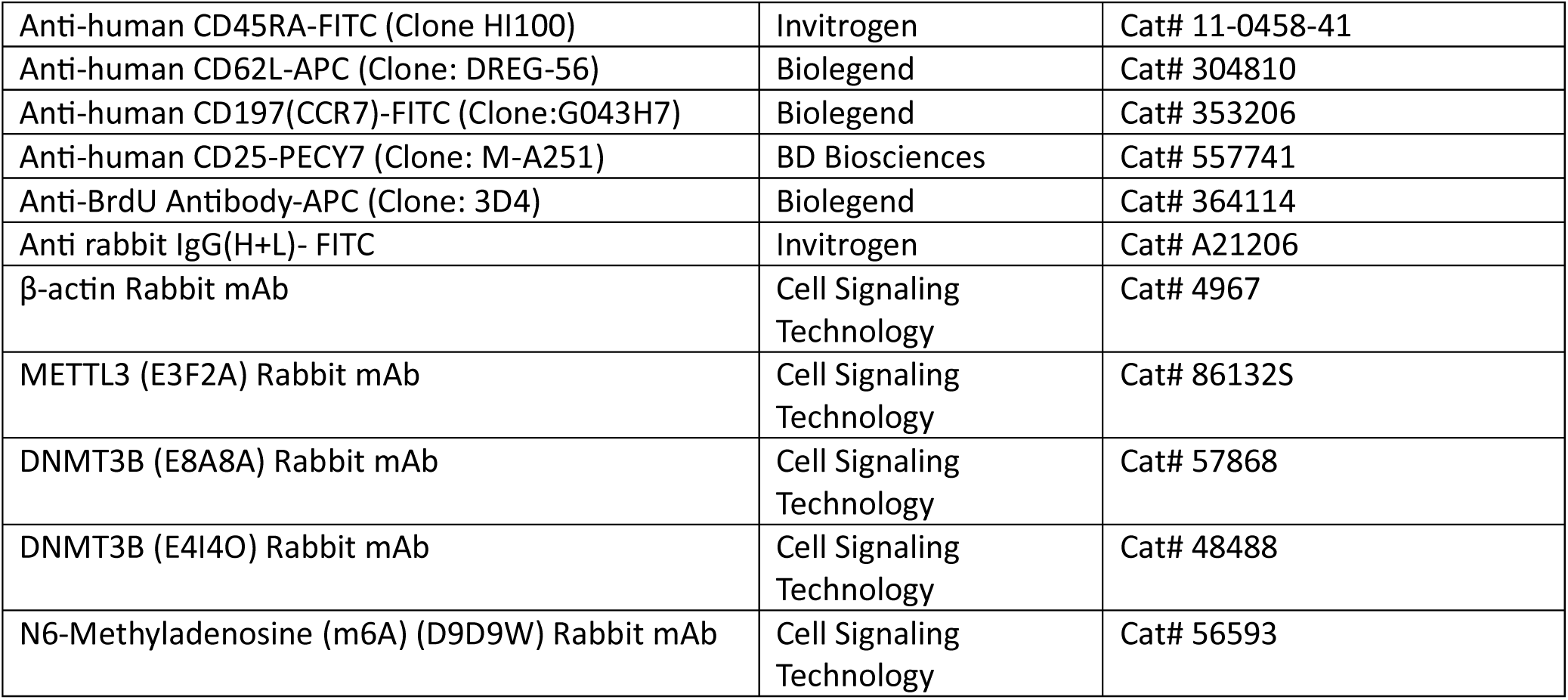
Antibodies.

**Table 3:** Instrument and software.

| Software/Algorithm | Source | Link |
| --- | --- | --- |
| FlowJo (v10) | BD Biosciences | https://www.flowjo.com/ |
| GraphPad Prism (v9) | GraphPad Software | https://www.graphpad.com/scientific-software/prism/ |
| Wave Software (Seahorse Analysis) | Agilent Technologies | https://www.agilent.com/ |
| ImageJ | NIH | https://imagej.net/ij |
| SRAMP (m6A prediction) | Zhou et al. | http://www.cuilab.cn/sramp |
| NCBI Primer-BLAST | NCBI | https://www.ncbi.nlm.nih.gov/tools/primer-blast/ |
| MethPrimer | Li & Dahiya | http://www.urogene.org/methprimer/ |
| R (v4.2.1) | R Core Team | https://www.r-project.org/ |
| R studio | R studio | Posit \| The Open-Source Data Science Company |

| Instrument/Equipment | Manufacturer |
| --- | --- |
| BD LSR Fortessa II Flow Cytometer | BD Biosciences |
| Agilent Seahorse XFe24 Analyzer | Agilent Technologies |
| Bio-Rad ChemiDoc Imaging System | Bio-Rad |
| CFX96 Real-Time PCR System | Bio-Rad |
| Multiskan SkyHigh Microplate Spectrophotometer | Thermo Fisher Scientific |
| Eppendorf Centrifuge 5810R | Eppendorf |
| T100 Thermal Cycler | Bio-Rad |
| Countess® II FL Automated Cell Counter | Thermo Fisher Scientific |

