## Supplementary figure legends and figures for "Mettl3-catalyzed m^6^A methylation determines CD8^+^ T cell differentiation fate in tumor"

**Figure S1. Characterization of CD8$^+$ T cells from murine tumors and in vitro models of T cell exhaustion.** (A) C57BL/6 mice were subcutaneously implanted with either MC38 or YUMM1.7, and CD8$^+$ T cells from the tumor-bearing host were assessed at the indicated times for the expression of PD1 vs. Tim3, and PD1 vs. CD39. Adjacent bar plots represent cumulative data from N=5 tumor-bearing mice. (B-C) UMAP embeddings of five integrated human tumor scRNA-seq profiles resolve (B) 14 transcriptionally distinct clusters, (C) with *CD8A* expression projected onto the same UMAP, identifying cluster 5 as intratumoral CD8$^+$ T cells. (D) Heatmap showing scaled expression of the top 10 differentially expressed genes across Tex, pTex1, and pTex2 clusters. (E) Module scores visualising enrichment of progenitor- and exhaustion-associated gene signatures across CD8$^+$ T cells. (F-J) Flow cytometry assessment of (F) PD1 vs. TIM3, (G) CTLA4, (H) TCF1 expression, and intracellular production of (I) IFNγ and (J) TNFα in CD8$^+$ T cells from healthy human donors either stimulated acutely (Acute) or expanded with chronic antigen stimulation (Chr). Adjacent bar plots show cumulative data from four independent experiments. *, $p < 0.05$; **, $p < 0.01$; ***, $p < 0.005$; ****, $p < 0.0001$.

**Figure S2. Mettl3 expression promotes exhaustion while impairing memory-like CD8$^+$ T cell differentiation. (**A) Western blot showing Mettl3 expression in naïve CD8$^+$ T cells and at different time points during in vitro activation. (B) Validation of Mettl3 knockdown using two independent lentiviral shRNAs (shMettl3 A and shMettl3 B) and overexpression (oeMettl3) using a lentiviral vector in human CD8$^+$ T cells. (C) CD8$^+$ T cells transduced with shMettl3 A, shMettl3 B, or an overexpression construct (oeMettl3) were expanded under Tex differentiation conditions and analyzed for PD1 vs. CD39 and PD1 vs. CTLA4 expression. The adjacent bar plot represents cumulative data from four independent experiments. (D-F) In vitro-differentiated $T_{Mem}$ cells from the indicated groups were analyzed for (D) FOXO1, (E)

CCR7, and (F) intracellular cytokine production (IFNγ and TNFα). Adjacent bar plots show cumulative data from four independent experiments. *, $p < 0.05$; **, $p < 0.01$; ***, $p < 0.005$; ****, $p < 0.0001$.

**Figure S3. Mettl3 confers an epigenetic bias that limits $T_{Mem}$ generation.** (A) Quantification of RNA $m^6A$ methylation levels in human $CD8^+$ T cells activated in the presence or absence of STM. (B-C) Expression of (B) CD25 and (C) CD44 in three-day-activated human $CD8^+$ T cells. Adjacent bar plots show cumulative data. (D) Extracellular acidification rate (ECAR) in response to glucose, oligomycin, and 2DG, with the adjacent bar plot showing glycolytic rate after glucose addition. (E) mRNA expression of glycolytic genes. (F) Oxygen consumption rate (OCR) over time in activated human $CD8^+$ T cells. Vertical lines indicate addition of mitochondrial inhibitors (oligomycin, FCCP, rotenone + antimycin A) for spare respiratory capacity (SRC) measurement; adjacent bar plot shows SRC ($OCR_{max}$ − OCR_basal). (G-H) Mean fluorescence intensity (MFI) of (G) MitoTracker Green and (H) MitoTracker Red in control and STM-treated $CD8^+$ T cells. (I) mRNA expression of mitochondrial fusion- and fission-related genes. (J) Proliferation of $CD8^+$ T cells (CTV-labeled) from healthy donors activated with or without STM. Representative plots show division peaks (P0-P5) with corresponding frequencies in control and STM-treated groups; adjacent bar graph summarizes frequencies across biological replicates. (K) Cell-cycle status determined by BrdU and 7AAD staining. Representative plots show percentages of apoptotic cells and cells in G0/G1, S, and G2/M phases; adjacent bar plots represent the distribution of these populations. Data are representative of six (A, G, and H), four (B-F, I-K) biological replicates. *, $p < 0.05$; **, $p < 0.01$; ***, $p < 0.005$; ****, $p < 0.0001$.

**Figure S4. The Mettl3-$m^6A$-Dnmt3b axis promotes epigenetic remodeling at memory-associated genes.** (A) $RT^2$ PCR Profiler Array data from two independent experiments showing fold changes in chromatin-modifying genes in $CD8^+$ T cells from healthy human donors activated for three days with or without STM. (B-C) Predicted $m^6A$ sites in *DNMT3B* mRNA using SRAMP: (B) confidence scores plotted across the transcript (vertical lines = site positions; dashed line = threshold); (C) table summarizing site positions, motif context, scoring metrics, and confidence classification. (D) *DNMT1* and *DNMT3A* mRNA decay analysis following Actinomycin D treatment in human $CD8^+$ T cells activated in the presence or absence of STM. Decay rates were determined by non-linear regression curve fitting (one-phase decay model). Data are representative of four independent experiments. (E) Dnmt3b expression in intratumoral $CD8^+$ T cells from MC38 and YUMM1.7 tumors. The adjacent bar plot represents cumulative data from six biological replicates. (F) Temporal dynamics of Dnmt3b expression in $CD8^+$ T cells differentiated into Tex or $T_{Mem}$. The adjacent bar graph shows cumulative data from six independent experiments. (G-I) Human $CD8^+$ T cells from the indicated groups differentiated into Tex and analyzed for: (G) PD1 and Tim3, (H) CTLA4, and (I) intracellular IFNγ and TNFα production. Adjacent bar graphs show cumulative data from four independent experiments. (J-M) Human $CD8^+$ T cells from the indicated groups differentiated into $T_{Mem}$ and analyzed for: (J) FOXO1, (K) TCF1, (L) CCR7, and (M) intracellular IFNγ and TNFα production. Adjacent bar graphs show cumulative data from five independent experiments. *, $p < 0.05$; **, $p < 0.01$; ***, $p < 0.005$; ****, $p < 0.0001$, ns=non-significant.

**Figure S5. Genetic knockdown of Mettl3 or Dnmt3b enhances $CD8^+$ T cell persistence and anti-tumor activity.** (A-B) Knockdown of (A) Mettl3 and (B) Dnmt3b in activated Pmel T cells using shMettl3 or shDnmt3b was validated by GFP expression (24 h post-lentiviral transduction) and western blotting. (C-F) Peripheral blood analysis from $Rag1^{-/-}$ mice at day 21 post-adoptive transfer with $Pmel^{Wt}$, $Pmel^{shMettl3}$, or $Pmel^{shDnmt3b}$ (prior to tumor rechallenge), showing frequencies of (C) central memory (CM; $CD44^+CD62L^+$) and effector memory (EM; $CD44^{+}CD62L^{-}$) subsets, (D) CCR7, (E) TCF1, and (F) FOXO1 expression in $CD8^{+}V\beta13^{+}$ T cells. (G-J) Peripheral blood analysis five days after B16F10 melanoma rechallenge in the same mice, showing (G) CM and EM percentages, (H) CCR7, (I) TCF1, and (J) FOXO1 expression in $CD8^{+}V\beta13^{+}$ T cells. (K-M) Tumor-infiltrating $CD8^{+}V\beta13^{+}$ T cells from rechallenged $Rag1^{-/-}$ mice were evaluated for (K) co-expression of PD1 with CTLA4 or CD39, and intracellular production of (L) IFNγ and (M) TNFα following re-stimulation. Adjacent bar plots (C-M) show cumulative data from *n* = 6 mice. (N-Q) $Pmel^{shMettl3}$ or $Pmel^{shDnmt3b}$ cells were adoptively transferred into tumor-bearing $Rag1^{-/-}$ mice. Tumor-draining lymph nodes (TdLN) and tumors were analyzed for (N, O) PD1/CTLA4 and (P, Q) PD1/CD39 expression in $V\beta13^{+}CD8^{+}$ T cells. Adjacent bar plots show cumulative data from *n* = 6 mice. *, $p < 0.05$; **, $p < 0.01$; ***, $p < 0.005$; ****, $p < 0.0001$.

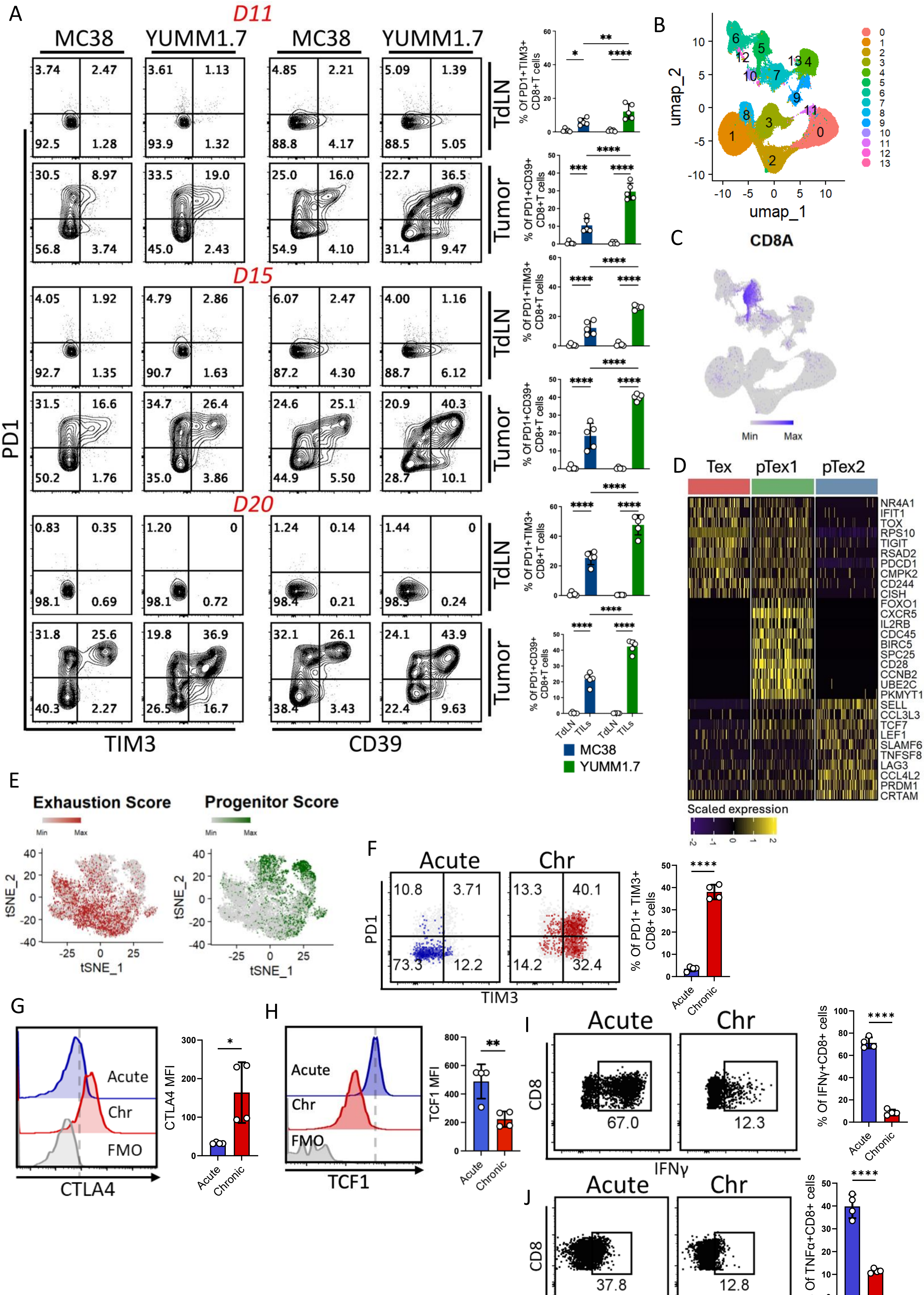
A
D11
MC38
YUMM1.7
TdLN
Tumor
D15
D20
PD1
TIM3
CD39
% Of PD1+TIM3+ CD8+T cells
% Of PD1+CD39+ CD8+T cells
TdLN
TILs
MC38
YUMM1.7
B
umap_2
umap_1
C
CD8A
Min
Max
D
Tex
pTex1
pTex2
NR4A1
IFIT1
TOX
RPS10
TIGIT
RSAD2
PDCD1
CMPK2
CD244
CISH
FOXO1
CXCR5
IL2RB
CDC45
BIRC5
SPC25
CD28
CCNB2
UBE2C
PKMYT1
SELL
CCL3L3
TCF7
LEF1
SLAMF6
TNFSF8
LAG3
CCL4L2
PRDM1
CRTAM
Scaled expression
E
Exhaustion Score
Progenitor Score
tSNE_2
tSNE_1
F
Acute
Chr
PD1
TIM3
% Of PD1+ TIM3+ CD8+ cells
Acute
Chronic
G
CTLA4
FMO
CTLA4 MFI
H
TCF1
TCF1 MFI
I
CD8
IFNγ
% Of IFNγ+CD8+ cells
J
TNFα
% Of TNFα+CD8+ cells

Figure S1

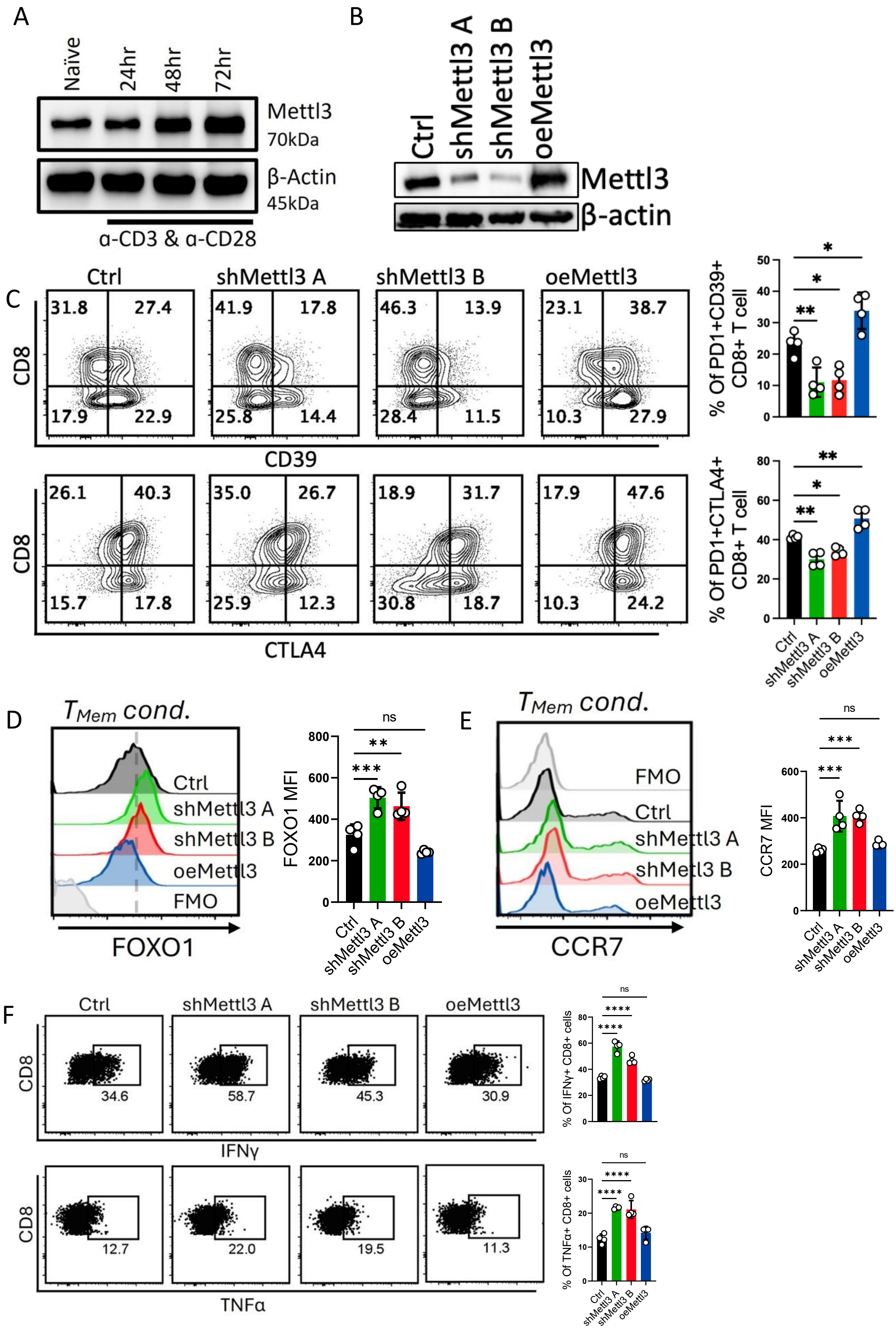
A
Naïve
24hr
48hr
72hr
Mettl3
70kDa
β-Actin
45kDa
α-CD3 & α-CD28
B
Ctrl
shMettl3 A
shMettl3 B
oeMettl3
Mettl3
β-actin
C
Ctrl
shMettl3 A
shMettl3 B
oeMettl3
CD8
CD39
CTLA4
% Of PD1+CD39+ CD8+ T cell
% Of PD1+CTLA4+ CD8+ T cell
D
TMem cond.
FOXO1
FOXO1 MFI
FMO
E
CCR7
CCR7 MFI
F
IFNγ
TNFα
% Of IFNγ+ CD8+ cells
% Of TNFα+ CD8+ cells

Figure S2

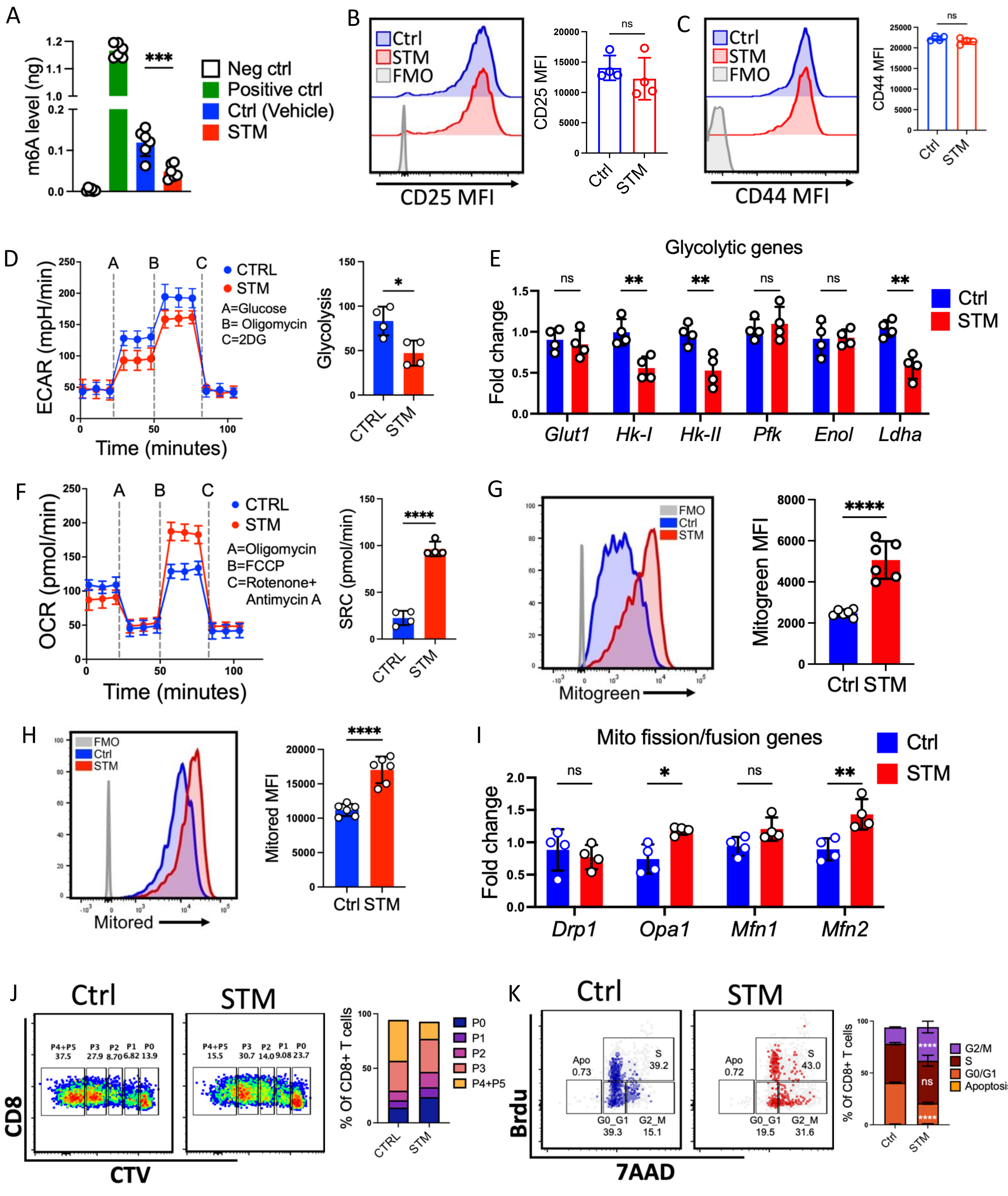
A
m6A level (ng)
***
Neg ctrl
Positive ctrl
Ctrl (Vehicle)
STM
B
Ctrl
STM
FMO
CD25 MFI
ns
C
CD44 MFI
D
ECAR (mpH/min)
Time (minutes)
CTRL
A=Glucose
B= Oligomycin
C=2DG
Glycolysis
E
Glycolytic genes
Fold change
Glut1
Hk-I
Hk-II
Pfk
Enol
Ldha
F
OCR (pmol/min)
A=Oligomycin
B=FCCP
C=Rotenone+
Antimycin A
SRC (pmol/min)
****
G
Mitogreen
Mitogreen MFI
H
Mitored
Mitored MFI
I
Mito fission/fusion genes
Drp1
Opa1
Mfn1
Mfn2
J
CD8
CTV
P4+P5
P3
P2
P1
P0
% Of CD8+ T cells
K
Apo
S
G0_G1
G2_M
Brdu
7AAD
G2/M
G0/G1
Apoptosis

Figure S3

A

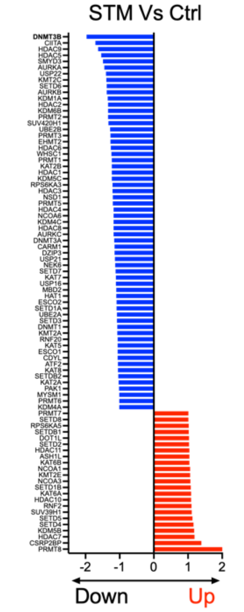


B

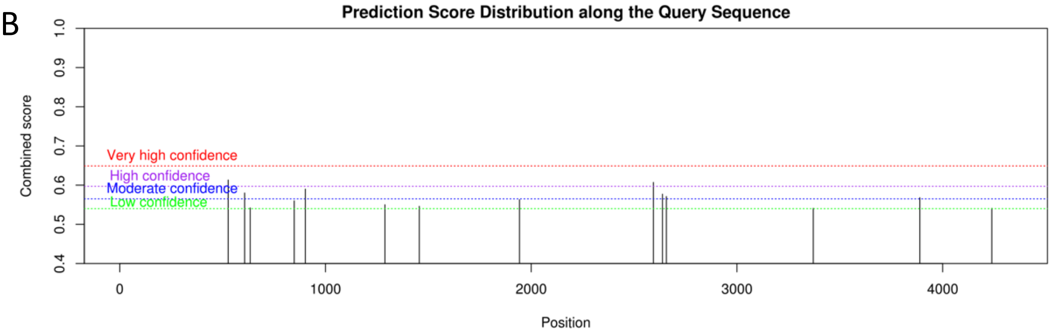


C

| Position | Sequence | Score(binary) | Score(knn) | Score(spectrum) | Score(combined) | Decision (m6A site) |
|---|---|---|---|---|---|---|
| 527 | AGUCUGCUAAGCUACACACAGGACUUGACAGGCGAUGGCGACGGG | 0.692 | 0.699 | 0.494 | 0.613 | High(H1) |
| 607 | AAAGCUCUUCCGGGAAACCAGGACUCGUUCAGAAAGCCCAGCUGU | 0.655 | 0.743 | 0.456 | 0.580 | Moderate(M1) |
| 634 | UUCAGAAAGCCCAGCUGUCCGAACUCGAAAUAACAACAGUGUCUC | 0.573 | 0.627 | 0.490 | 0.542 | Low(L1) |
| 848 | GACCUCACAGACGACACAGAGGACACACAUGGGACGCCCCAGAGC | 0.581 | 0.717 | 0.511 | 0.560 | Low(L2) |
| 902 | CCCUACGCCCGCCUAGCCCAGGACAGCCAGCAGGGGGGCAUGGAG | 0.545 | 0.644 | 0.646 | 0.590 | Moderate(M2) |
| 1289 | ACCUUCCCCAGCAGCCCUGGAGACUCAUUGGAGGACCAGCUGAAG | 0.581 | 0.610 | 0.500 | 0.550 | Low(L3) |
| 1456 | AUCAAGGAAAUACGAGAACAAGACUCGAAGACGCACAGCUGACGA | 0.583 | 0.645 | 0.483 | 0.546 | Low(L4) |
| 1943 | GGCGUCCUGCGGCGCCGGAAGGACUGGAACGUGCGCCUGCAGGCC | 0.656 | 0.800 | 0.406 | 0.563 | Low(L5) |
| 2594 | AAUGAUAAACUCGAGCUGCAGGACUGCUUGGAAUACAAUAGGAUA | 0.679 | 0.643 | 0.504 | 0.607 | High(H2) |
| 2638 | AGCCAAGUUAAAGAAAGUACAGACAAUAACCACCAAGUCGAACUC | 0.588 | 0.602 | 0.559 | 0.577 | Moderate(M3) |
| 2657 | CAGACAAUAACCACCAAGUCGAACUCGAUCAAACAGGGGAAAAAC | 0.557 | 0.403 | 0.612 | 0.571 | Moderate(M4) |
| 3371 | GUUGUAGGCAUGUAGCUACAGGACAUUUUUAAGGGCCCAGGAUCG | 0.645 | 0.317 | 0.426 | 0.541 | Low(L6) |
| 3889 | GAGGCAGUGACAGCAGUCAGGGACAGACAUACAUUUCUCAUACCU | 0.596 | 0.466 | 0.543 | 0.568 | Moderate(M5) |
| 4239 | ACGUAACAAAUAUGGGGAAAAAACUGUGCCUUGUUUCAACAGUUU | 0.570 | 0.277 | 0.535 | 0.541 | Low(L7) |

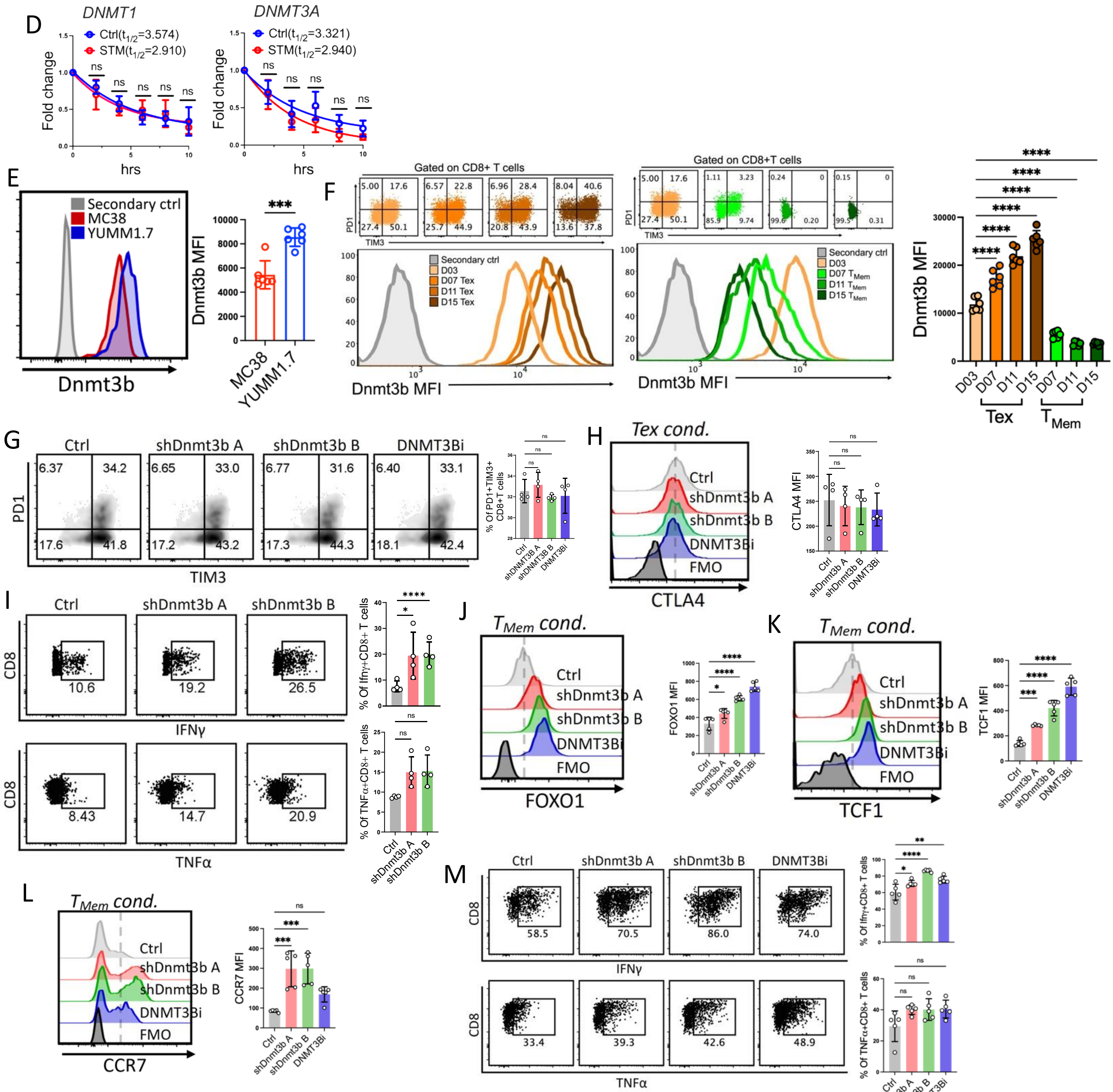


Figure S4

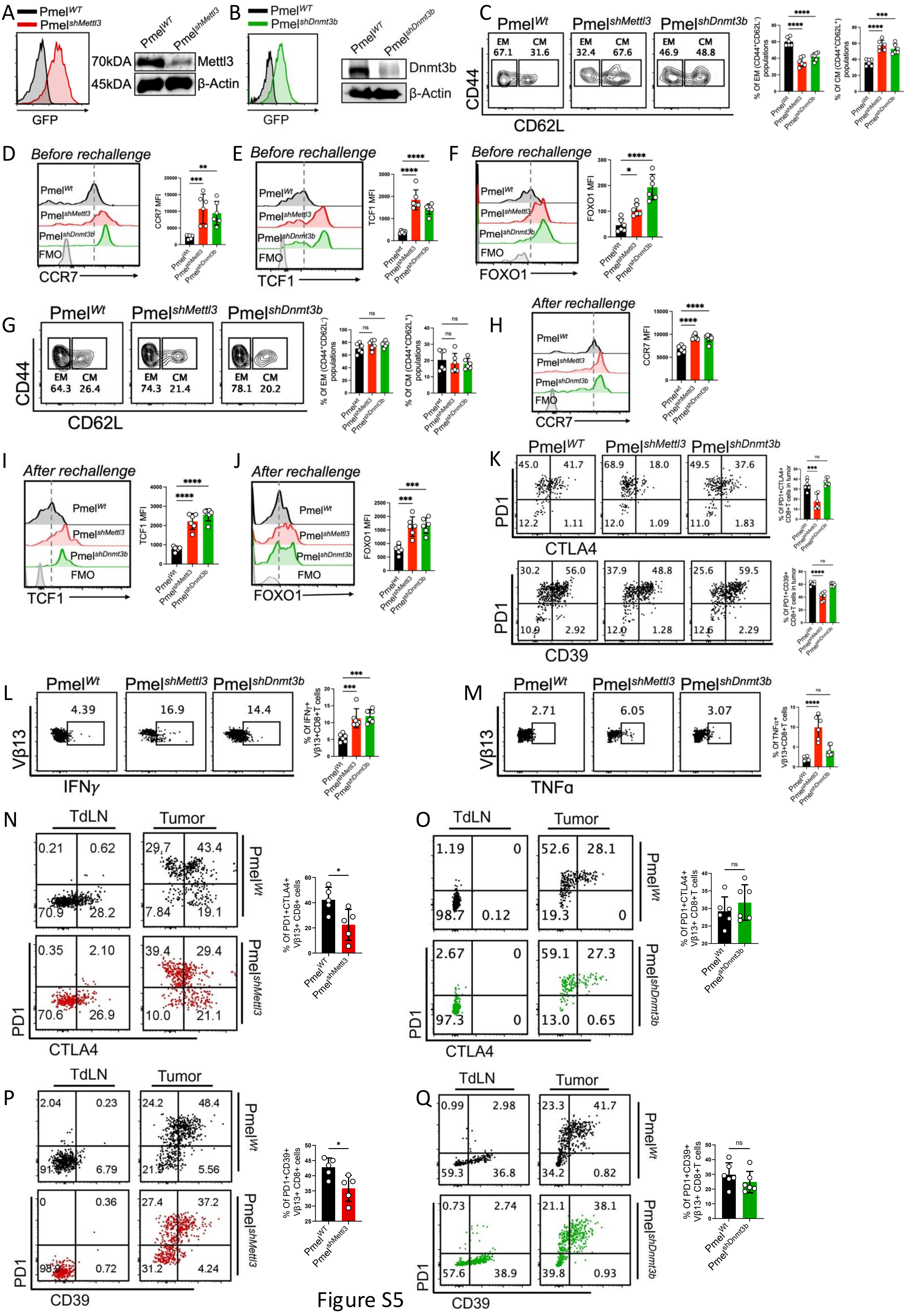
A
Pmel^WT
Pmel^shMettl3
GFP
70kDA
45kDA
Mettl3
β-Actin
B
Pmel^shDnmt3b
Dnmt3b
C
CD44
CD62L
EM
CM
% Of EM (CD44+CD62L-) populations
% Of CM (CD44+CD62L+) populations
D
Before rechallenge
CCR7
FMO
CCR7 MFI
E
TCF1
TCF1 MFI
F
FOXO1
FOXO1 MFI
G
H
After rechallenge
I
J
K
PD1
CTLA4
CD39
% Of PD1+CTLA4+ CD8+T cells in tumor
% Of PD1+CD39+ CD8+T cells in tumor
L
Vβ13
IFNγ
% Of IFNγ+ Vβ13+CD8+T cells
M
TNFα
% Of TNFα+ Vβ13+CD8+T cells
N
TdLN
Tumor
% Of PD1+CTLA4+ Vβ13+ CD8+ cells
O
% Of PD1+CTLA4+ Vβ13+ CD8+T cells
P
% Of PD1+CD39+ Vβ13+ CD8+ cells
Q
% Of PD1+CD39+ Vβ13+ CD8+T cells

Figure S5
